# Identification of basic helix-loop-helix transcription factors that activate betulinic acid biosynthesis by RNA-sequencing of hydroponically cultured *Lotus japonicus*

**DOI:** 10.1101/2022.11.16.516519

**Authors:** Hayato Suzuki, Hirokazu Takahashi, Ery Odette Fukushima, Mikio Nakazono, Toshiya Muranaka, Hikaru Seki

## Abstract

Although triterpenes are ubiquitous in plant kingdom, their biosynthetic regulatory mechanisms are limitedly understood. Here, we found that hydroponic culture dramatically activated betulinic acid (BA) biosynthesis in the model Fabaceae *Lotus japonicus*, and investigated its transcriptional regulation. Fabaceae plants develop secondary aerenchyma (SA) on the surface of hypocotyls and roots during flooding for root air diffusion. Hydroponic culture induced SA in *L. japonicus* and simultaneously increased the accumulation of BA and the transcript levels of its biosynthetic genes. RNA-sequencing of soil-grown and hydroponically cultured plant tissues, including SA collected by laser microdissection, revealed that several transcription factor genes were co-upregulated with BA biosynthetic genes. Overexpression of *LjbHLH32* and *LjbHLH50* in *L. japonicus* transgenic hairy roots upregulated the expression of BA biosynthesis genes, resulting in enhanced BA accumulation. However, transient luciferase reporter assays in *Arabidopsis* mesophyll cell protoplasts showed that LjbHLH32 transactivated promoters of biosynthetic genes in the soyasaponin pathway but not the BA pathway, like its homolog GubHLH3, a soyasaponin biosynthesis regulator in *Glycyrrhiza uralensis*. This suggested the evolutionary origin and complex regulatory mechanisms of BA biosynthesis in Fabaceae. This study sheds light on the unrevealed biosynthetic regulatory mechanisms of triterpenes in Fabaceae plants.

**Highlight:** Hydroponic culture enhanced betulinic acid synthesis in *Lotus japonicus*. RNA-sequencing and functional characterization experiments suggest that LjbHLH32 and LjbHLH50 are the transcription factors activating betulinic acid biosynthesis.

## Introduction

Triterpenes are plant specialized (secondary) metabolites with diverse structures and biological properties (Thimmappa *et al*., 2014). The common precursor, 2,3-oxidosqualene, is composed of six isoprene units supplied by the mevalonate (MVA) pathway. Multiple oxidosqualene cyclases (OSCs), cytochrome P450 monooxygenases (P450s), and UDP-dependent glycosyltransferases (UGTs) catalyze conversion of the substrates to structurally diversified non-glycosylated triterpenes and triterpene glycosides (saponins; Seki *et al*., 2015).

Fabaceae plants produce a large amount of triterpenes with marked chemodiversity. Soyasaponins are specific to Fabaceae plants such as *Glycine max*, *Medicago* spp., *Glycyrrhiza* spp., and *Lotus japonicus* (Fenwick, 1992; Hayashi *et al*., 1998; Tsuno *et al*., 2018). *Medicago truncatula* produces oleanolic acid (OA)-derived hemolytic saponins (Huhman *et al*., 2005). Thickened roots of *Glycyrrhiza uralensis* accumulate glycyrrhizin, but more OA and betulinic acid (BA) were produced in tissue-cultured stolons (Kojoma *et al*., 2010; Tamura *et al*., 2017). *L. japonicus* accumulates C-28-oxidized triterpenes [ursolic acid (UA), OA, and BA] in a tissue-dependent manner (Suzuki *et al*., 2019). Biosynthesis pathways of these Fabaceae triterpenes have been almost understood by the recent identification of CSyGT (cellulose-synthase-derived glycosyltransferase) enzymes catalyzing the glucuronosylation of triterpene aglycones (Jozwiak *et al*., 2020; Chung *et al*., 2020). However, their physiological functions and biosynthetic regulatory mechanisms are unclear.

Plant genomes possess a large number of basic helix-loop-helix (bHLH)-type transcription factors (TFs), which are classified into approximately 25 clades based on sequence similarity in the bHLH domain and other conserved domains (Heim *et al*., 2003). Fabaceae plants possess more clade IVa bHLH genes than related plant families, which are subdivided into three subclades (IVa1-3) based on a phylogenetic tree of the full-length proteins (Suzuki *et al*., 2021). Some members of subclade IVa1 are regulators of Fabaceae triterpene saponin biosynthesis. TSAR1 and TSAR2 are positive regulators of soyasaponins and hemolytic saponins, respectively, in *M. truncatula* (Mertens *et al*., 2016). TSAR3, their paralog, is a regulator of hemolytic saponin biosynthesis during seed development (Ribeiro *et al*., 2020). GubHLH3, a TSAR homolog, upregulated soyasaponin biosynthesis in *G. uralensis* (Tamura *et al*., 2018). TFs regulating BA biosynthesis have yet been identified in Fabaceae plants. On the other hand, BpbHLH9 (clade Ia) was identified as an OA and BA biosynthesis regulator in the Betulaceae plant *Betula platyphylla* (Yin *et al*., 2017). The clade Ib bHLH TFs Bl and Bt are regulators of cucurbitane-type triterpenes in leaves and fruits, respectively, of cucumber (Cucurbitaceae; Shang *et al*., 2014). Thus, identification of triterpene-related bHLH TFs in *L. japonicus* provides insight into the evolutionary history of triterpene biosynthesis regulators in the Fabaceae and other plant families.

We performed triterpene profiling to characterize triterpene production in *L. japonicus*. A large amount of BA accumulated in secondary aerenchyma (SA) tissue in hypocotyl and roots when cultured hydroponically. SA is a white, spongy tissue developed from phellogen [secondary meristem (SM)], whose formation is an adaptive trait of Fabaceae plants to survive flooding by promoting oxygen diffusion in roots (Mochizuki *et al*., 2000; Yamauchi *et al*., 2013). Because the expression of BA biosynthetic genes (*OSC3* and *CYP716A51*) was upregulated in whole roots of hydroponically cultured plants and SA, we performed comparative RNA sequencing (RNA-seq) analysis to identify candidate TFs that positively regulate BA biosynthesis. From comparisons between soil-cultured and hydroponically cultured whole roots and between SA and SM collected by laser microdissection (LMD), we identified 23 TFs co-upregulated in hydroponic roots and SA. Among them, we focused on LjbHLH32 (subclade IVa2) and LjbHLH50 (clade Ia) based on their co-expression with BA biosynthetic genes and phylogenetic analysis. The functions of these two bHLH-type TFs in triterpene biosynthesis regulation were examined and are discussed.

## Materials and Methods

### Plant materials

Seeds of *L. japonicus* Gifu B-129 accession and *M. truncatula* cv. R108 and Jemalong A17 were sterilized using a five-fold-diluted sodium hypochlorite solution containing 0.02% Tween-20 for 15 min with mild agitation. After washing three times in sterilized water, they were soaked in water with mild agitation overnight. The seeds were germinated on 0.8% agar with incubation for 4 days (*L. japonicus*) or 3 days (*M. truncatula*) in the dark and for 2 days on a 16 h light/8 h dark cycle at 25°C. The seedlings were transplanted to soil pots or hydroponic solutions and cultured in a plant growth room maintained at 23°C until harvest. We used basal nutrient solution (BNS) for all hydroponic solutions (Conn *et al*., 2013). *L. japonicus* seedlings were cultured in 5 mL sampling tubes for 2 weeks and transferred to 50 mL sampling tubes with BNS. *M. truncatula* seedlings were cultured in 50 mL tubes with BNS from transplant to harvest. For induction of adventitious roots, the roots of 1-week-old seedlings were isolated and cultured in 100 mL of half-strength Gamborg’s B5 liquid medium supplemented with 3% sucrose and 0.2 mg/L of indole-3-butyric acid. The culture was maintained at 23°C with shaking at 90 rpm and subcultured every 3 to 4 weeks. The *L. japonicus* Lj suspension cell line (rpc00032) was maintained at 27°C with shaking at 120 rpm in Murashige and Skoog (MS) medium (pH 5.7) supplemented with 1 mg/L of 2,4-D and 3% sucrose, and subcultured every 7 days. For metabolite analysis of cultured cells, ethanol or 100 μM methyl jasmonate (MeJA) was added and the cells were cultured for 4 additional days.

### Triterpene extraction and gas chromatography-mass spectrometry (GC-MS) analysis

Triterpenes were extracted from 10 mg of dried plant powder and analyzed by GC-MS following our previous report (Suzuki *et al*., 2019).

### RNA extraction from whole roots

Whole roots (including the hypocotyl) of 2.5-month-old *L. japonicus* cultured on soil or in hydroponics were harvested and immediately frozen in liquid nitrogen. Total RNA extraction, recombinant DNase I treatment, and column purification were performed as described previously (Suzuki *et al*., 2019).

### RNA extraction from SA and SM

Sample fixation, paraffin-embedding, microtome sectioning, collection of SA and SM cell layers by LMD, and RNA extraction were performed following a previous report (Takahashi *et al*., 2010) with some modifications.

Hypocotyls of *L. japonicus* seedlings cultured hydroponically for 7 weeks were harvested. Segments of hypocotyls were obtained at 5 to 10 mm from the cotyledons. The 5-mm-long hypocotyls were fixed in methanol by incubation for 30 min under a vacuum of 0.09 MPa. This procedure was repeated three times with new methanol.

We employed a microwave-assisted method for paraffin-embedding using an Energy Beam Sciences H2850 Microwave Tissue Processor (Agawan, MA). A vial containing fixed tissue and pre-chilled methanol was placed on ice and microwaved three times at 250 W at 37°C for 15 min. The methanol was replaced each time. The temperature in the vial was < 20°C during the first and second microwave treatments; however, 37°C was allowed in the third treatment. The solvent was gradually replaced with 1-butanol, using methanol:1-butanol (3:1), methanol:1-butanol (1:1), methanol:1-butanol (1:3), and 1-butanol, with microwaving at each step at 300 W and 37°C for 1.5 min. 1-Butanol was gradually replaced with melted paraffin wax (Surgipath Paraplast X-tra Tissue Infiltration/Embedding Media; Leica Biosystems, Wetzlar, Germany) using 1-butanol:paraffin (3:1), 1-butanol:paraffin (1:1), and 1-butanol:paraffin (1:3) with microwaving at each step at 300 W and 58°C for 10 min. Two treatments at 250 W at 58°C for 10 min and five treatments at 250 W at 58°C for 30 min were performed with new paraffin solution. The specimens were embedded in paraffin wax and cooled to room temperature. The paraffin blocks were stored at 4°C.

RNAsecure reagent (Thermo Fisher Scientific, Waltham, MA) was diluted 25-fold with UltraPure Distilled Water (Thermo Fisher Scientific). The diluted RNAsecure solution was placed on an Arcturus PEN Membrane Glass Slide (Thermo Fisher Scientific) heated on a hot plate (57°C). Paraffin sections (12 μm thickness) were floated on the surface of the solution and incubated for several tens of seconds. After removing the solution, the sections were dried at 42°C for 1 h. The slides were immersed in Histoclear II (National Diagnostics, Atlanta, GA) for 10 min twice to remove the paraffin, and air-dried. LMD was performed using a PALM MicroBeam IV Laser-capture Microdissection System (Zeiss, Jena, Germany) with a MicroBeam Laser (cut energy: 55–58, cut focus: 80, LPC energy: 30, LPC focus: 2). AdhesiveCap 500 clear (Zeiss) was used to collect cell layers cut by the laser. RNA was purified using an Arcturus PicoPure RNA Isolation Kit (Thermo Fisher Scientific).

### Quantitative reverse transcription-PCR (qRT-PCR) analysis

First-strand cDNA was synthesized from purified total RNA using PrimeScript RT Master Mix (Perfect Real Time; Takara Bio, Shiga, Japan) following the manufacturer’s instructions. The reaction time was 15 and 60 min, for total RNA of whole roots and that of LMD samples, respectively. We performed qRT-PCR analysis following a previous report (Suzuki *et al*., 2019) using primer nos. 1–20 in Table S1.

### RNA-seq

Transcriptome profiling of soil-grown and hydroponic roots was performed with three biological replicates (Table S2). Complementary DNA library construction was performed using a TruSeq Stranded mRNA Library Prep Kit (Illumina, San Diego, CA) according to the manufacturer’s instructions. The resulting cDNA libraries were sequenced using HiSeq 2500 (Illumina) with 100 bp single-end (SE) reads. Adapter sequences and low-quality reads were removed from raw RNA-seq reads by Trimmomatic v. 0.38 (Bolger *et al*., 2014). Quantification of transcripts was conducted with Salmon v. 0.14.2 (Patro *et al*., 2017). Reference *L. japonicus* coding sequences (predicted from genome assembly build v. 1.2 of Gifu B-129 accession; Kamal *et al*., 2020) were downloaded from *Lotus* Base (https://lotus.au.dk/). A mapping-based index was built from the reference using an auxiliary k-mer hash over k-mers of length 31 as recommended. Summarized gene counts were obtained from the transcript counts using the R package tximport v. 1.16.1 (Soneson *et al*., 2015). Differentially expressed genes (DEGs) were identified using the graphical user interface for TCC (TCC-GUI) (Su *et al*., 2019). Low count genes (0 mapped reads) were filtered out from TCC computation and default settings were kept for the other parameters.

Transcriptome profiling of the LMD samples was performed with four biological replicates. RNA-seq libraries were constructed and sequenced using a Novaseq 6000 (Illumina) with 150-bp paired-end (PE) reads by Takara Bio. Because two of four replicates of SA samples had smaller total read numbers, they were combined into one FastQ file and analyzed with three biological replicates (Table S2). Data processing was conducted as described above after removing unpaired reads.

### Gene cloning

The primers used are listed in Table S1. PCR was performed using Prime STAR MAX or Prime STAR GXL DNA Polymerase (Takara Bio). The plasmid pENTR1A (Thermo Fisher Scientific) was linearized by PCR (primer nos. 21 and 22). The full coding sequences of *LjbHLH32* and *LjbHLH50* were amplified from cDNA of hydroponically cultured roots using primer nos. 24–27 and were integrated into pENTR1A using NEBuilder HiFi DNA Assembly Master Mix (New England Biolabs, Ipswich, MA) to obtain gateway entry clones (pENTR1A-*LjbHLH32* and pENTR1A-*LjbHLH50*). *GFP* was amplified from pUB-GWS-GFP (GenBank accession no. AB303066) using primer nos. 28 and 29, and cloned into pENTR1A to obtain pENTR1A-*GFP*.

### Construction of the gateway destination vector pSD11

The binary vector pUB-GW-GFP is used for overexpression in *L. japonicus* hairy roots (Maekawa *et al*., 2008). After constructing pUB-GW-GFP, the *Arabidopsis thaliana* heat shock protein (HSP) 18.2 terminator, which is suitable for gene overexpression in plants, was identified (Nagaya *et al*., 2010; Matsui *et al*., 2014). Here, we constructed the binary vector pSD11 by replacing the terminators of pUB-GW-GFP with the HSP terminator (Fig. S1).

The primers used are listed in Table S1. The 875 bp native *AtHSP18.2* terminator (HSPter) was amplified from *A. thaliana* Col-0 genomic DNA using primer nos. 30 and 31, and cloned into pJET1.2 using a CloneJET PCR Cloning Kit (Thermo Fisher Scientific). The vector backbone was amplified from pUB-GWS-GFP using primer nos. 32 and 33. The HSP terminator was amplified from pJET1.2-HSPter using primer nos. 34 and 35. These two fragments were assembled by NEBuilder HiFi DNA Assembly Master Mix (New England Biolabs) to construct pSD8.

The vector backbone, including the gateway cassette and GFP, and fragments, including the *LjUBQ1* promoter, were amplified from pUB-GWS-GFP using primer sets 33/36 and 38/39, respectively. These two fragments were assembled with the HSP terminator fragment (primer nos. 34 and 37) to produce pSD9.

The gateway destination vector pSD11 was constructed from PCR-amplified fragments of pSD8 (primer nos. 40 and 41) and pSD9 (primer nos. 42 and 43) by the golden gate cloning method using BsaI-HF v. 2 and T4 DNA ligase (both New England Biolabs).

### Generation and analysis of TF-overexpressing hairy roots

The expression vectors (pSD11-*GFP*, *LjbHLH32*, and *LjbHLH50*) were generated by LR reaction of pSD11 and entry clones (pENTR1A-*GFP*, *LjbHLH32*, and *LjbHLH50*) using Gateway LR Clonase II Enzyme Mix (Thermo Fisher Scientific). These three vectors were introduced into *Agrobacterium rhizogenes* ATCC15834 by electroporation.

*Agrobacterium rhizogenes*-mediated hairy root transformation was performed following our previous study (Suzuki *et al*., 2019), with some modifications. The incubation temperature was maintained at 25°C in the germination, co-culture, and hairy root induction steps. Isolated hairy roots were cultured at 23°C. For liquid culture of hairy roots, the rotary shaker agitation speed was set to 90 rpm. The germination/co-cultivation medium contained 1/2 B5 basal salts. Hairy root elongation (HRE) medium is composed of 1/2 B5 basal salts, 1/4 Gamborg’s B5 vitamins, and 300 mg/L of cefotaxime sodium. All media were adjusted to pH 5.5 before autoclaving and freshly prepared. Seeds were germinated as described above. After infection and co-cultivation (Suzuki *et al*., 2019), seedlings were placed on HRE agar supplemented with 1% sucrose and a filter paper strip was placed on the hypocotyls to prevent the hairy roots from drying. Hairy roots were elongated under a 16 h light/8 h dark cycle for 3 weeks, followed by shoot removal. Isolated hypocotyls possessing hairy roots were incubated on new HRE agar containing 3% sucrose without light for 10 days. Single hairy roots were isolated and cultured in 5 mL of liquid HRE medium supplemented with 3% sucrose using six-well cell culture plates. They were subcultured twice every 16 days. Hairy root lines emitting GFP fluorescence were selected and each was subcultured into six wells. After 26 days of culture, hairy roots were harvested for GC-MS, qRT-PCR, and RNA-seq analysis (150 bp PE reads, Novaseq 6000 platform, by Takara Bio; Table S2) as described above.

### Cloning of promoter sequences

Putative promoter sequences, including the 5’-untranslated region (2–2.5 kb upstream from the start codon), of triterpene biosynthetic genes were retrieved using the sequence retrieval tool in *Lotus* Base (https://lotus.au.dk/; Table S3). The primers used to amplify the promoter sequences are listed in Table S1 (primer nos. 44–67). The plasmid pENTR1A (Thermo Fisher Scientific) was linearized by PCR (primer nos. 21 and 23). The promoter sequences of *OSC1*, *OSC3*, *OSC9*, *OSC5*, *CYP716A51*, *LjCYP93E1*, *LjCYP72A61*, *LjCSyGT*, putative 2,3-dihydro-3,5-dihydroxy-6-methyl-4*H*-pyran-4-one (DDMP) transferase (*LjDDMPT*), HMG-CoA reductase 1 (*LjHMGR1*), and squalene epoxidase 1 (*LjSQE1*) were integrated into pENTR1A by NEBuilder HiFi DNA assembly cloning.

### Vector construction for transient effector-reporter analysis

The *Sfi*I-digested PCR fragment, including the gateway cassette (primer nos. 68 and 69 in Table S1) and *Sfi*I-digested pGL4.1HSP [encoding firefly luciferase (fLUC); Nakata *et al*., 2013], were ligated using T4 DNA ligase (New England Biolabs) to generate the pGL4.1HSP-GW vector. Promoter sequences were integrated into pGL4.1HSP-GW by the LR reaction to generate 11 reporter constructs (pGL4.1HSP-GW-*OSC1pro*, *OSC3pro*, *OSC9pro*, *OSC5pro*, *CYP716A51pro*, *LjCYP93E1pro*, *LjCYP72A61pro*, *LjCSyGTpro*, *LjDDMPTpro*, *LjHMGR1pro*, and *LjSQE1pro*). pENTR-D-*GuCYP93E3pro* (Tamura *et al*., 2018) was used to generate pGL4.1HSP-GW-*GuCYP93E3pro*.

TF genes were transferred from entry clones [pENTR-D-*GubHLH3* (Tamura *et al*., 2018), pENTR1A-*LjbHLH32*, and pENTR1A-*LjbHLH50*] to pDEST35SHSP (Oshima *et al*., 2013) to produce three effector constructs (pDEST35SHSP-*GubHLH3*, *LjbHLH32*, and *LjbHLH50*). The plasmid pDEST35SHSP-*vesicle-associated membrane protein 722* (*Vamp722*; Yoshida *et al*., 2013) was used as a negative control.

We employed phRLHSP [encoding a modified *Renilla* luciferase (RLUC); Yoshida *et al*., 2013] as an internal reference to normalize the results of our dual luciferase reporter assays.

### Transient effector-reporter analysis

*A. thaliana* Col-0 was grown under a 16 h light/8 h dark cycle at 22°C for 3–4 weeks. Rosette leaves were used for protoplast isolation after peeling by the Tape-*Arabidopsis* Sandwich method (Wu *et al*., 2009). Protoplasts were isolated as described (Yoshida *et al*., 2013) with minor modifications. Peeled leaves were incubated for 1 h in enzyme solution [400 mM mannitol, 20 mM KCl, 10 mM CaCl_2_, 10 g/L of cellulase onozuka R10 (Yakult Pharmaceutical Industry Co., Ltd., Tokyo, Japan), 2.5 g/L of macerozyme R10 (Yakult Pharmaceutical Industry Co., Ltd.), 10 mM 2-mercaptoethanol, and 20 mM 2-morpholinoethanesulfonic acid (MES), pH 5.7] in a shaker (22°C, 60 rpm). The protoplasts were filtered through a 70 μm cell strainer and collected by centrifugation at 100 × *g* for 10 min. The supernatant was removed by decantation and the remaining pellet was washed thrice with W5 buffer [150 mM NaCl, 125 mM CaCl_2_, 5 mM KCl, and 2 mM MES, pH 5.7] followed by centrifugation at 100 × *g* for 5 min. The pellet was washed with MMg solution [400 mM mannitol, 15 mM MgCl_2_, and 4 mM MES, pH 5.7] and resuspended in 5 mL of MMg solution (when using 24 leaves).

Reporter, effector, and reference plasmids were co-transfected into protoplasts by the polyethylene glycol (PEG)-assisted method (Yoshida *et al*., 2013). After incubating transfected protoplasts at 22°C for 16–19 h, FLUC and RLUC activities were measured using a Pikka Gene Dual Assay Kit (Toyo Ink, Tokyo, Japan).

### Phylogenetic analysis

Sequences were collected as described previously (Suzuki *et al*., 2021). The full-length peptide sequence of CjbHLH1 (BAJ40865.1; Yamada *et al*., 2011) was retrieved from GenBank (https://www.ncbi.nlm.nih.gov/genbank/). The sequences of Bl (Csa5g156220) and Bt (Csa5g157230; Shang *et al*., 2014) were obtained from the Cucurbit Genomics Database (http://cucurbitgenomics.org/). A phylogenetic tree was constructed by the neighbor-joining method with 1000 replicates using MEGA6 software (Tamura *et al*., 2013) following alignment by MUSCLE (Edgar, 2004).

A phylogenetic tree of lateral organ boundaries domain (LBD) proteins was generated using Clustal Omega v. 1.2.3 (Sievers *et al*., 2011), FastTree v. 2.1.10 (Price *et al*., 2010), and MEGA X (Kumar *et al*., 2018) following our previous report (Suzuki *et al*., 2021).

## Results

### Hydroponic culture enhances BA accumulation in *L. japonicus* hypocotyl and roots

Previously, we reported triterpene profiling in the model Fabaceae *L. japonicus* grown on soil (Fig. S2; Suzuki *et al*., 2019). To investigate triterpene production characteristics in this plant, we conducted triterpene sapogenin profiling of whole roots from soil-grown and hydroponically cultured plants, and liquid-cultured adventitious roots (Figs. 1A and S3). Because triterpenes and saponins were extracted and acid-hydrolyzed to remove sugar moieties attached to aglycones, we detected soyasapogenols instead of soyasaponins by GC-MS. BA (peak 6) and soyasapogenol B (SB; a major soyasapogenol in this plant, peak 10) were detected in the extract of soil-grown roots. Liquid culture of adventitious roots induced by indole-3-butyric acid accumulated BA and SB with lower peak intensities. Hydroponic roots of *L. japonicus* accumulated a larger amount of BA than soil-cultured roots and adventitious roots, which caused peak leading in the GC-MS chromatogram. BA was detected in the extracts of roots without acid hydrolysis, indicating that BA is not glycosylated *in planta*. MeJA induces soyasaponin but not BA biosynthesis in Fabaceae plants (Hayashi *et al*., 2003; Broeckling *et al*., 2005; Suzuki *et al*., 2005). We analyzed Lj suspension cultured cells with and without MeJA (Figs. 1B and S3). Neither BA nor SB was detected in the extract of untreated cultured cells. Extracts of the MeJA-treated cells contained a small amount of SB but no BA.

**Fig. 1.**
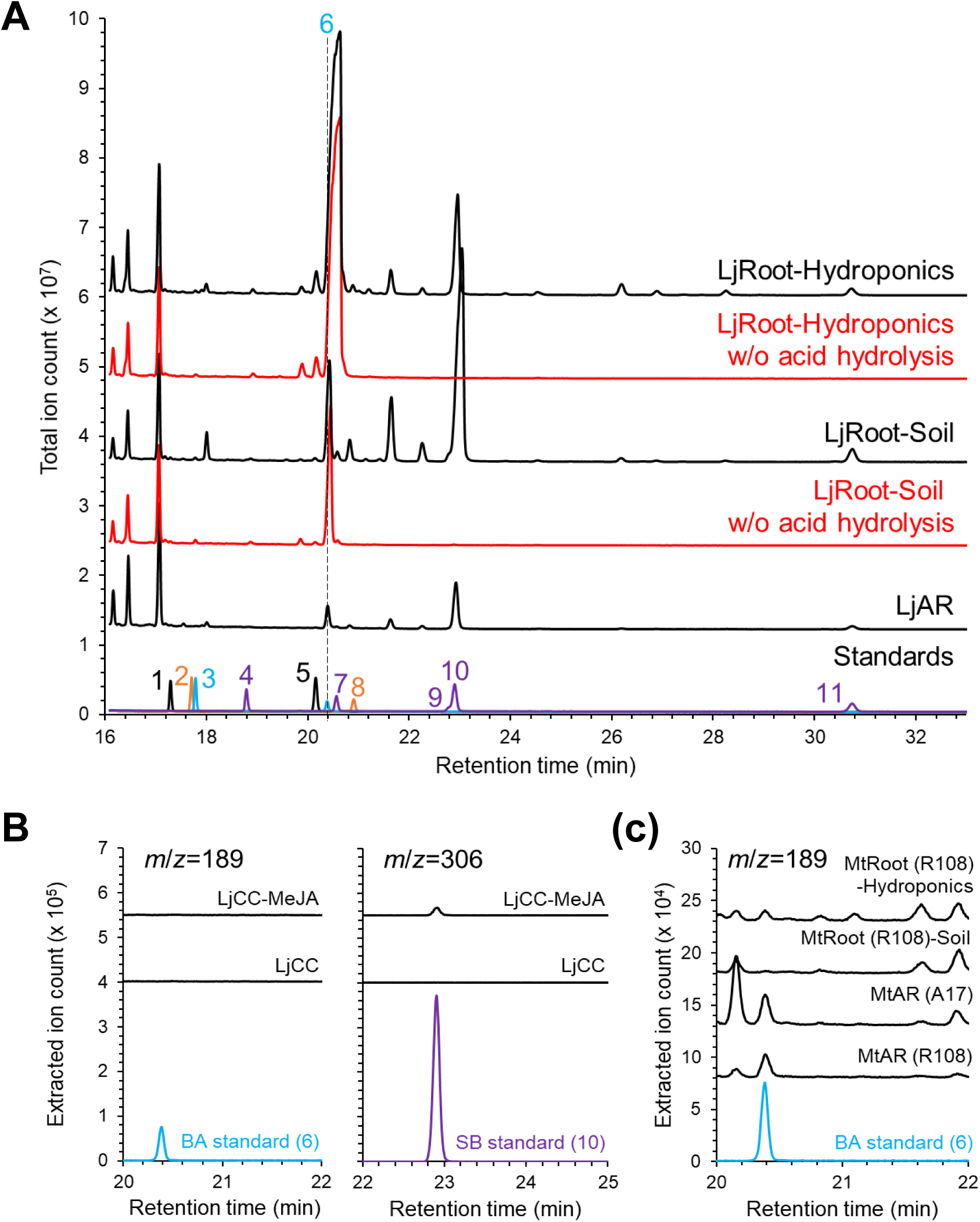
Triterpene sapogenin profiling by gas chromatography-mass spectrometry (GC-MS). **(A)** Analysis of *Lotus japonicus* adventitious roots (AR) and whole roots of soil-grown and hydroponically cultured plants. Increased betulinic acid (BA) accumulation was observed in the extracts of hydroponic roots. A BA peak was observed without acid hydrolysis during triterpene extraction. **(B)** Analysis of Lj suspension cultured cells (LjCC) with and without methyl jasmonate (MeJA). Extracted ion chromatograms (*m/z* = 189 and 306) are shown to assess the presence of BA and soyasapogenol B (SB). **(C)** Extracted ion chromatograms (*m/z* = 189) of hydroponically cultured and soil-grown roots and AR of *Medicago truncatula* cv. R108 and Jemalong A17. The following authentic standards were analyzed (Suzuki *et al*., 2019): β-amyrin (1), α-amyrin (2), lupeol (3), 24-OH-β-amyrin (4), oleanolic acid (5), BA (6), sophoradiol (7), ursolic acid (8), soyasapogenol E (9), SB (10), soyasapogenol A (11).

We also analyzed the model Fabaceae plant *M. truncatula* (Figs. 1C and S3). Although soil-grown roots of *M. truncatula* (cv. R108) did not accumulate BA, BA was detected in the liquid-cultured adventitious roots and roots of hydroponically cultured plants. To our knowledge, this is the first report of BA accumulation in *Medicago* spp. We also analyzed the adventitious roots of cv. Jemalong A17 and confirmed the presence of BA. However, the amount of BA in *M. truncatula* was much lower than that in *L. japonicus*. *L. japonicus* is more appropriate to study the regulatory mechanisms of BA biosynthesis enhanced in hydroponic culture.

### SA produces and accumulates BA

Next, we evaluated BA production in hydroponic roots. We analyzed triterpenes extracted from whole roots of *L. japonicus* plants hydroponically cultured for 2–12 weeks and observed a time course-dependent increase in BA and enlargement of the hypocotyl and main roots (Fig. S4A–C). Development of a white, spongy tissue was confirmed by anatomical analysis (Fig. 2A), and we assumed that this tissue was homologous to SA in *G. max* (soybean). Soybean develops SA from the SM on stem, hypocotyl, main and lateral roots, and nodules as an adaptive trait to survive flooding by promoting oxygen diffusion in roots (Shimamura *et al*., 2010; Yamauchi *et al*., 2013). Hydroponic culture likely resulted in a lack of oxygen in roots, similar to flooding. Hydroponic roots accumulated 10-fold more BA than soil-grown roots, and its quantity reached 6.5 mg/100 mg-dry weight (Fig. 2B). Isolated SA accumulated a higher amount of BA (> 12 mg/100 mg-dry weight) than stele and lateral roots (Fig. 2B). However, SB did not increase in hydroponic roots and SA (Fig. 2C).

**Fig. 2.**
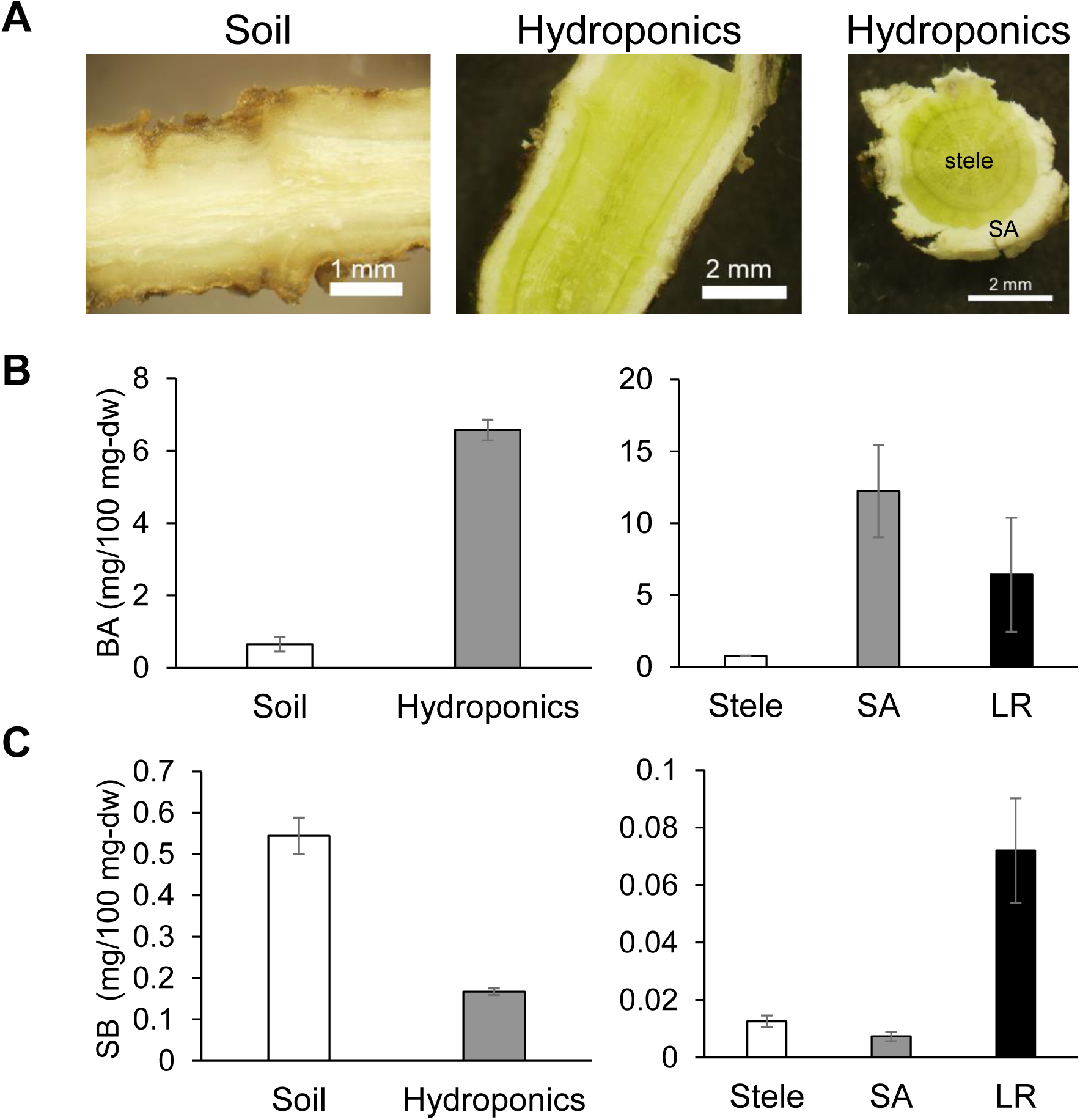
Accumulation of betulinic acid (BA) in secondary aerenchyma (SA) tissue induced by hydroponic culture. **(A)** Images of sections of hypocotyls. White, spongy tissue (SA) had developed on the surface of hypocotyls and roots of *L. japonicus* cultured hydroponically. Quantitation of BA **(B)** and soyasapogenol B (SB; **C)**. Hydroponic roots and SA accumulated more BA than soil-grown roots (Soil) and stele, respectively. Means ± standard errors (SEs, n = 3). LR, lateral roots.

We analyzed the expression levels of biosynthetic genes for BA (*OSC3* and *CYP716A51*) and SB (*OSC1* and *CYP93E1*) (Fig. 3) by qRT-PCR. When comparing soil-grown and hydroponic roots, BA biosynthetic genes were upregulated in hydroponic roots (Fig. 3). For metabolite analysis (Fig. 2), we harvested SA, including cortical cells. To identify the tissue biosynthesizing BA, we isolated SA and SM cell layers by LMD (Fig. S5) and performed RNA extraction and qRT-PCR. Because BA biosynthetic gene expression was higher in SA (Fig. 3), more BA is likely synthesized in SA. These results suggest that transcriptional regulation is responsible for the increased BA level in SA of *L. japonicus* hydroponic roots.

**Fig. 3.**
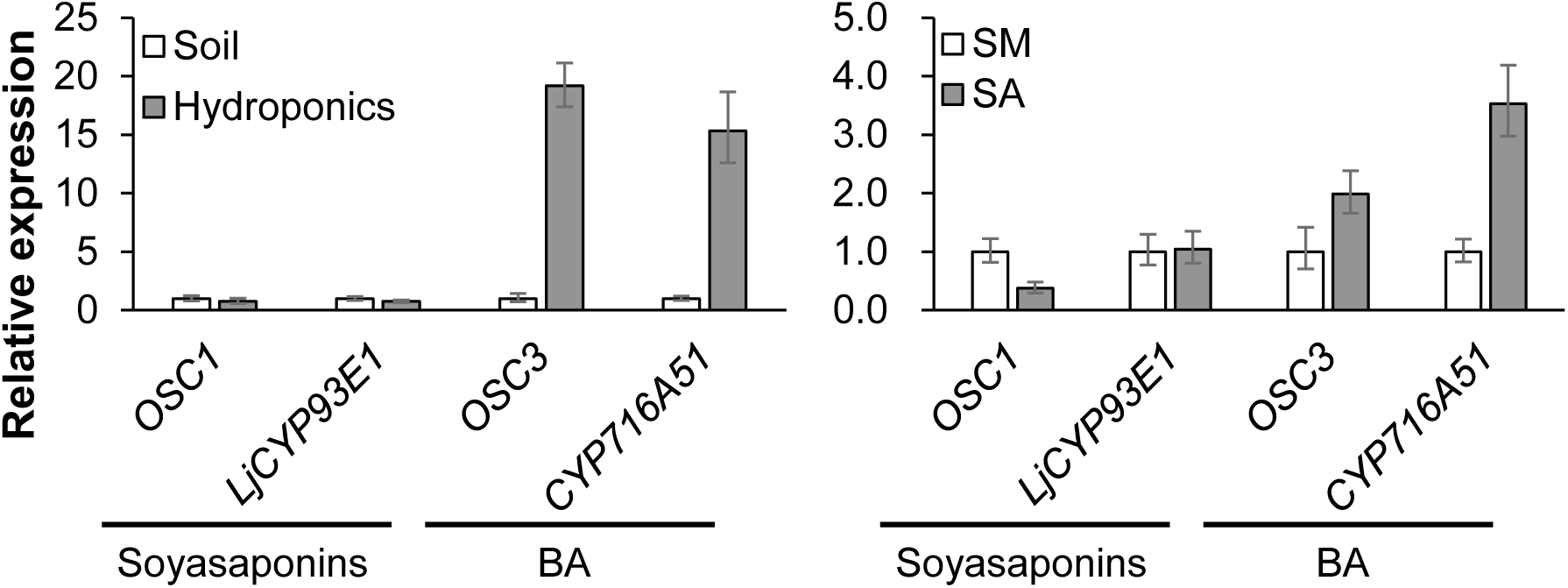
Quantitative reverse transcription-PCR (qRT-PCR) analysis of triterpene biosynthetic genes. Relative expression levels of betulinic acid (BA) and soyasaponin biosynthesis genes were measured in triplicate. The values were normalized to the expression level of a reference gene (*LjUBQ1*). Means ± standard deviations (SDs). Soil, soil-grown roots; hydroponics, hydroponically grown roots; SM, secondary meristem; secondary aerenchyma, SA.

### Candidate regulators of BA biosynthesis as identified by RNA-seq

We performed an RNA-seq-based comparative transcriptome analysis for mining potential transcriptional regulators of BA biosynthesis. RNA evaluated by qRT-PCR (Fig. 3) was used for the analysis, and sequencing of the whole root samples and the LMD samples was conducted in three and four replicates, respectively. Data processing was performed in triplicate for all samples as mentioned in Materials and Methods. We performed differential expression analyses independently in data set 1 [soil-grown roots (So) and hydroponic roots (Hy)] (Figs. 4A and S6A) and data set 2 (SM and SA) (Figs. 4B and S6B). Based on the MA plot (Fig. S6), we identified 3152 and 1994 DEGs whose expression was higher in So and Hy, respectively (Table S4). SM and SA had 2169 and 1647 DEGs with elevated expression levels (Table S4). There were 256 DEGs overlapping between Hy and SA (Fig. 4C), including 23 putative TF genes (Fig. 4C and Table S5), as well as BA biosynthetic genes (*OSC3* and *CYP716A51*) and genes for rate-limiting enzymes for triterpene biosynthesis [3-hydroxy-3-methyl-glutaryl-CoA reductase1 (HMGR1) and squalene epoxidase1 (SQE1)].

**Fig. 4.**
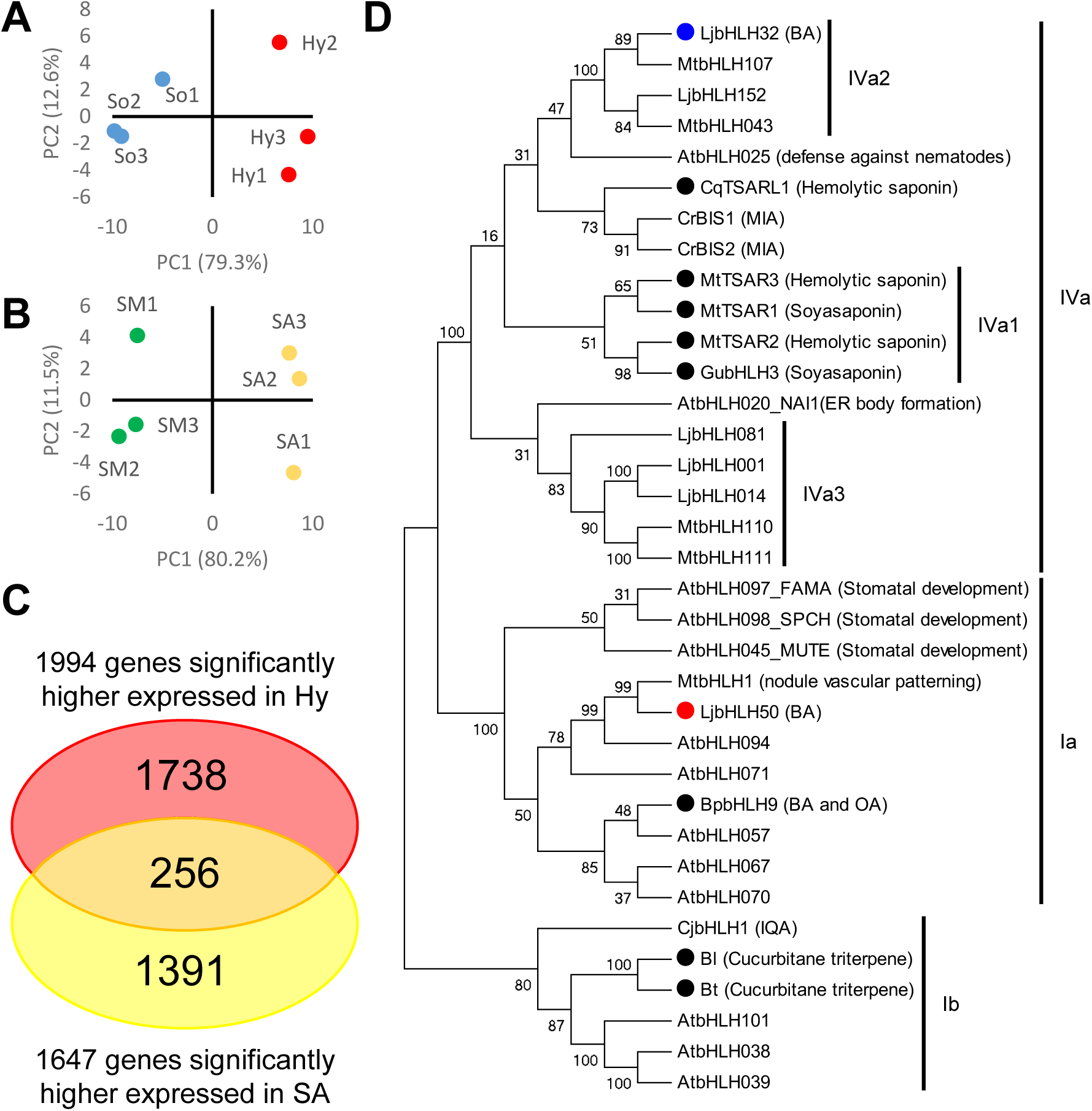
Selection of candidate transcription factors (TFs) regulating betulinic acid (BA) biosynthesis. Principal component analysis (PCA) of the transcriptomes of soil-grown (So) and hydroponically cultured (Hy) roots **(B)**, and secondary meristem (SM) and secondary aerenchyma (SA) tissues **(B)**. **(C)** Venn diagram of differentially expressed genes (DEGs) upregulated in Hy and SA compared to So and SM, respectively. **(D)** Phylogenetic relationships of basic helix-loop-helix proteins. TFs known to regulate triterpene biosynthesis are highlighted by black circles. Blue and red circles indicate LjbHLH32 and LjbHLH50, respectively. ER, endoplasmic reticulum; OA, oleanolic acid; IQA, isoquinoline alkaloid.

We assessed the correlation between the expression patterns of BA biosynthetic genes and those of candidate TF genes using the gene expression atlas in *Lotus* Base (https://lotus.au.dk/expat/; 81 conditions based on annotation and prediction in Miyakojima MG-20 genome v. 3.0). Seven TF genes were co-expressed with *OSC3*, *CYP716A51*, *HMGR1*, and *SQE1* (Pearson’s correlation coefficient [PCC] threshold: 0.4; Table 1). Among the seven TFs, we selected LjbHLH32 (LotjaGi4g1v0043000) and LjbHLH50 (LotjaGi1g1v0340900) for further analysis because bHLH TFs are triterpene biosynthesis regulators in Fabaceae (Mertens *et al*., 2016; Tamura *et al*., 2018), *Panax* spp. (Zhang *et al*., 2017), cucumber (Shang *et al*., 2014), and *B. platyphylla* (Yin *et al*., 2017).

**Table 1.** Co-expression analysis of triterpene biosynthetic genes and 23 candidate TF genes.

TF genes
| Name | Gene ID |  | Pearson's correlation coefficient* |  |  |  |
| --- | --- | --- | --- | --- | --- | --- |
|  | Gifu B-129<br>ver 1.2 | Miyakojima<br>MG-20<br>ver 3.0 | *OSC3* | *HMGR1* | *CYP716A51* | *SQE1* |
| OSC3 | LotjaGi2g1v0235400 | Lj2g3v0914630 | 1.00 | 0.51 | 0.49 | 0.50 |
| HMGR1 | LotjaGi2g1v0352700 | Lj2g3v1574250 | 0.51 | 1.00 | 0.66 | 0.68 |
| CYP716A51 | LotjaGi4g1v0438900 | Lj4g3v3014670 | 0.49 | 0.66 | 1.00 | 0.63 |
| SQE1 | LotjaGi5g1v0037500 | Lj5g3v0347610 | 0.50 | 0.68 | 0.63 | 1.00 |
| NAC | LotjaGi1g1v0222300 | Lj0g3v0171479 | 0.20 | 0.24 | 0.25 | 0.31 |
| **bHLH50 (Ia)** | **LotjaGi1g1v0340900** | **Lj1g3v1525930** | **0.57** | **0.68** | **0.58** | **0.85** |
| MYB | LotjaGi1g1v0548000 | - | - | - | - | - |
| MYB | LotjaGi1g1v0650400 | Lj1g3v4012910 | -0.54 | -0.47 | -0.54 | -0.75 |
| bHLH (Ib) | LotjaGi1g1v0667300 | Lj0g3v0202379 | 0.23 | 0.14 | 0.07 | 0.48 |
| WRKY | LotjaGi1g1v0694700 | Lj1g3v4578840 | -0.13 | -0.03 | -0.21 | -0.25 |
| **bZIP** | **LotjaGi1g1v0724300** | **Lj1g3v4315340** | **0.44** | **0.62** | **0.44** | **0.54** |
| **WRKY** | **LotjaGi1g1v0754400** | **Lj1g3v4830200** | **0.63** | **0.77** | **0.54** | **0.64** |
| Zinc finger | LotjaGi2g1v0209200 | Lj2g3v0740620 | - | - | - | - |
| **LBD** | **LotjaGi2g1v0242300** | **Lj2g3v0934070** | **0.58** | **0.79** | **0.81** | **0.82** |
| ERF | LotjaGi3g1v0009300 | Lj0g3v0243159 | -0.28 | -0.48 | -0.27 | -0.39 |
| LBD | LotjaGi3g1v0035700 | Lj3g3v0397200 | -0.01 | 0.30 | 0.67 | 0.26 |
| SCREAM-like protein | LotjaGi3g1v0203400 | Lj3g3v1404080 | - | - | - | - |
| MYB_related | LotjaGi3g1v0380600 | Lj0g3v0318769 | -0.29 | -0.24 | -0.07 | -0.11 |
| NAC | LotjaGi4g1v0002900 | Lj4g3v0050650 | -0.49 | -0.34 | -0.31 | -0.55 |
| **bHLH32 (IVa2)** | **LotjaGi4g1v0043000** | **Lj0g3v0292969** | **0.74** | **0.79** | **0.85** | **0.67** |
| **WRKY** | **LotjaGi4g1v0379800** | **Lj4g3v2616130** | **0.67** | **0.70** | **0.47** | **0.69** |
| ERF | LotjaGi4g1v0422400 | Lj4g3v2871880 | 0.39 | 0.50 | 0.34 | 0.56 |
| **Zinc finger** | **LotjaGi5g1v0010400** | **Lj5g3v0165540** | **0.66** | **0.66** | **0.56** | **0.82** |
| NAC | LotjaGi5g1v0158000 | Lj5g3v1093960 | 0.38 | 0.39 | 0.25 | 0.22 |
| Zinc finger | LotjaGi5g1v0173100 | - | - | - | - | - |
| LBD | LotjaGi5g1v0323400 | Lj5g3v2099760 | 0.10 | 0.09 | 0.10 | 0.44 |
| bHLH (IVc) | LotjaGi6g1v0344900 | Lj6g3v2168640 | -0.38 | -0.43 | -0.39 | -0.52 |

\*Pearson's correlation coefficient (PCC) values based on transcriptome data of MG-20 deposited in
*Lotus* Base. Bold indicates putative TF genes showing a > 0.4 of PCC to four BA biosynthetic genes.

The bHLH TFs were classified into approximately 25 clades based on sequence similarity of the bHLH domain and other conserved domains (Heim *et al*., 2003). We constructed a phylogenetic tree of full-length bHLH proteins of clades Ia, Ib, and IVa, including representative bHLH proteins involved in specialized plant metabolism (Fig. 4D). Fabaceae clade IVa bHLH TFs were further subdivided into three subclades, and subclade IVa1 included all Fabaceae triterpene saponin biosynthesis activators identified in clade IVa (Suzuki *et al*., 2021). Interestingly, LjbHLH32 was a member of subclade IVa2 but not IVa1 (Fig. 4D). The orthologs in subclade IVa2 were highly conserved in Fabaceae plants and were commonly expressed in roots and nodules of model Fabaceae plants (Suzuki *et al*., 2021). However, no subclade IVa2 bHLHs have been functionally characterized to date. The closest homolog of another candidate LjbHLH50 in *M. truncatula* is MtbHLH1 (Medtr3g099620), which is suggested to regulate nitrogen transfer between nodule and root cells by controlling nodule vascular patterning (Godiard *et al*., 2011). LjbHLH50 and MtbHLH1 showed 60% amino acid identity (99% coverage) and belonged to clade Ia (Fig. 4D), similar to BpbHLH9, an OA and BA biosynthesis regulator in *B. platyphylla* (Yin *et al*., 2017). These phylogenetic relationships between our candidates and known triterpene biosynthesis regulators suggest that LjbHLH32 and LjbHLH50 are involved in BA biosynthesis regulation in *L. japonicus*.

### Overexpression of LjbHLH32 or LjbHLH50 enhances BA production in transgenic hairy roots

We generated GFP-expressing (control), LjbHLH32-overexpressing (LjbHLH32-OX), and LjbHLH50-OX hairy root lines for functional identification of LjbHLH32 and LjbHLH50 *in planta*. We chose three independent lines for each construct and analyzed their triterpene content by GC-MS (Fig. 5A). Compared to the control lines, LjbHLH32-OX and LjbHLH50-OX hairy roots accumulated 5.73- and 4.31-fold more BA (Fig. 5A, right), but the amount of SB was weakly elevated (Fig. 5A, left). Next, we assessed the expression of triterpene biosynthetic genes by qRT-PCR (Fig. 5B). LjbHLHs-OX lines overexpressed the corresponding bHLH genes. LjbHLH32 overexpression increased the transcript levels of the BA biosynthetic genes *LjHMGR1*, *LjSQE1*, *OSC3*, and *CYP716A51*, whereas branch pathway genes [cycloartenol synthase gene (*OSC5*), *OSC1*, and *LjCYP93E1*] were only slightly affected. LjbHLH50-OX lines showed 1.56–2.26-fold higher expression of *LjHMGR1*, *LjSQE1*, *OSC5*, *OSC1*, and *LjCYP93E1*. However, the expression of *OSC3* and *CYP716A51* was enhanced to a greater degree.

**Fig. 5.**
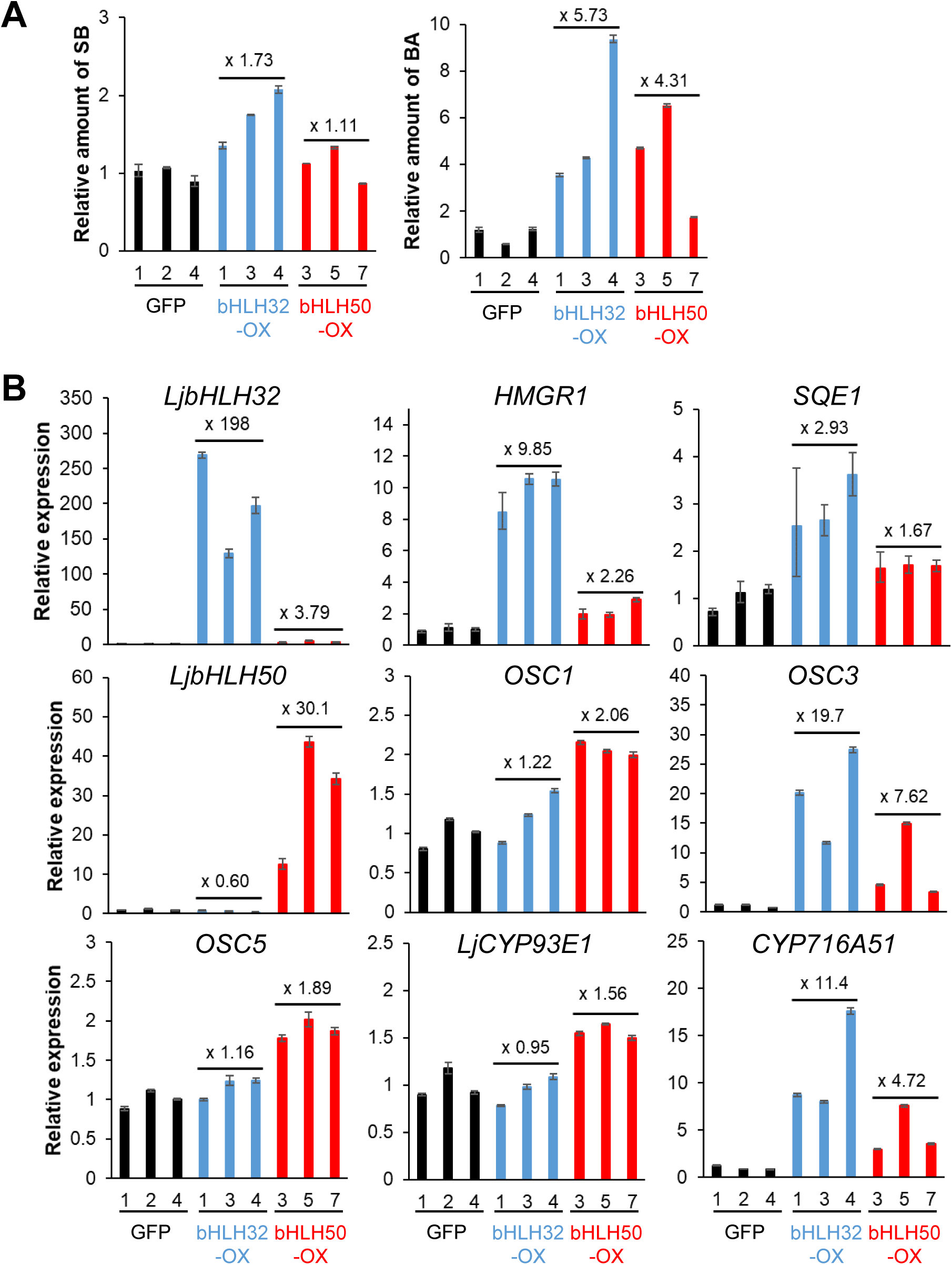
Metabolite and gene expression analysis of LjbHLH32- and LjbHLH50-overexpressing (OX) transgenic hairy root lines. **(A)** Triterpenes were extracted from three control (GFP-expressing) lines and a basic helix-loop-helix (bHLH)-OX line. Accumulation of betulinic acid (BA) and soyasapogenol B (SB) in hairy roots was analyzed by gas chromatography-mass spectrometry (GC-MS). Relative amounts were calculated from the peak areas of BA and SB. **(B)** Relative expression levels of bHLH and biosynthetic genes by quantitative reverse transcription-PCR (qRT-PCR). Error bars indicate standard deviations (SD, n = 3).

We performed an RNA-seq analysis of nine hairy root lines (Fig. 5) to test the effect of LjbHLH32 and LjbHLH50 overexpression on a genome-wide scale. The control lines and bHLH-OX lines were distinguished by principal component analysis (PCA) of transcriptome data (Fig. 6A and B). DEGs were identified based on an MA plot (Fig. S6C and D). Compared to the control lines, 527 and 1045 genes were significantly upregulated in LjbHLH32-OX and LjbHLH50-OX, respectively (Table S4). Among the DEGs upregulated in bHLH-OX hairy roots and 256 DEGs commonly upregulated in Hy and SA samples (Fig. 4C), 6 overlapped (Fig. 6C). These six genes included the BA biosynthetic genes *OSC3* and *CYP716A51* but no genes involved in other triterpene biosynthesis pathways (Table S6). Some of the enzyme genes putatively involved in precursor supply for BA biosynthesis were significantly upregulated by overexpression of bHLHs (Table S7), which may contribute to increased BA production in hairy roots (Fig. 5A).

**Fig. 6.**
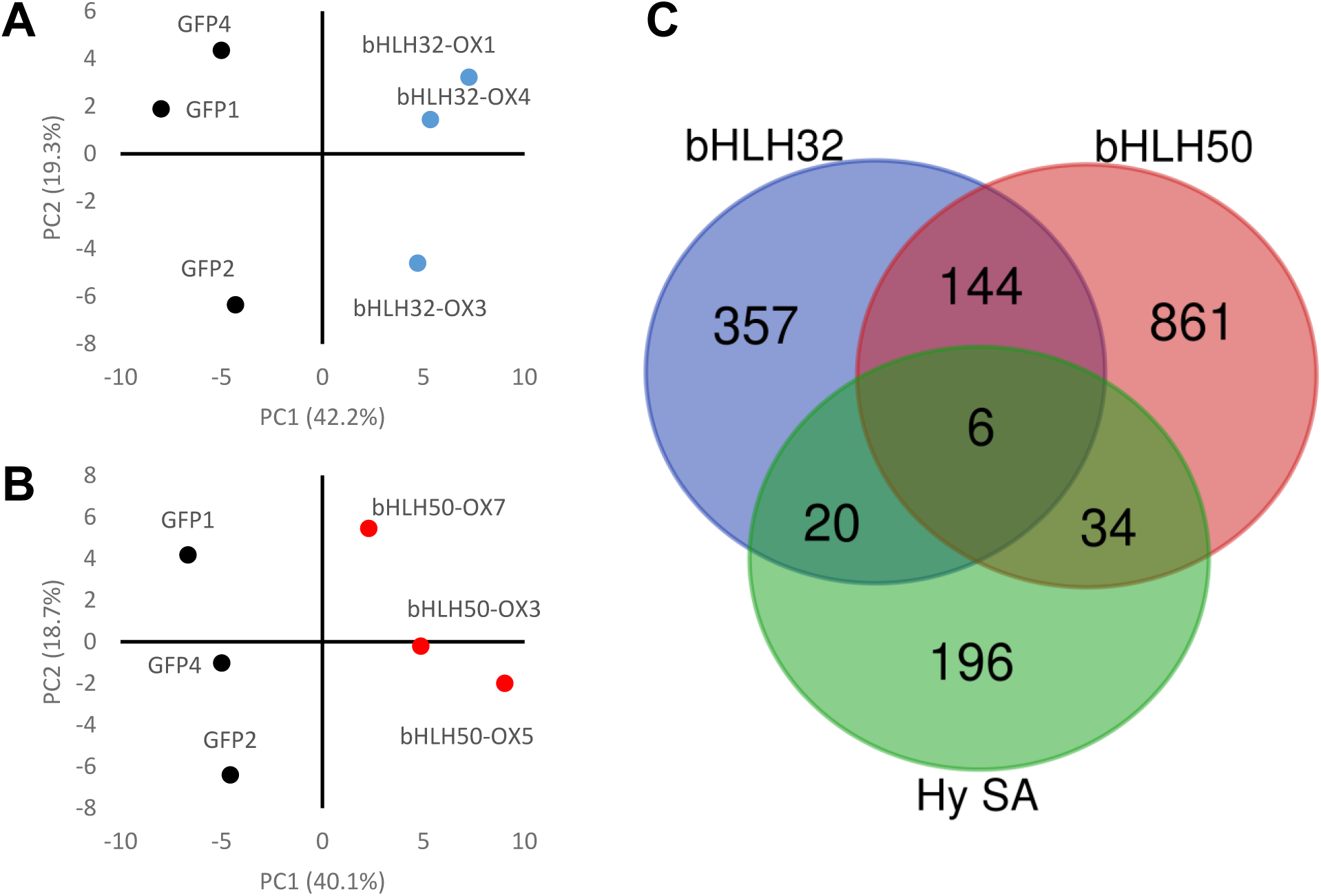
RNA-sequencing (RNA-seq) analysis of LjbHLH32- and LjbHLH50-overexpressing (OX) transgenic hairy root lines. Principal component analysis (PCA) of the transcriptomes of control and LjbHLH32-OX roots **(A)**, and control and LjbHLH50-OX roots **(B)**. **(C)** Venn diagram of 256 differentially expressed genes (DEGs) commonly upregulated in hydroponic whole roots (Hy) and secondary aerenchyma (SA), and DEGs upregulated in bHLH-OX hairy root lines compared to control (GFP-expressing) hairy roots.

The genes upregulated in the LjbHLH32- and LjbHLH50-OX lines (Fig. 6C) are likely involved in the metabolism of primary metabolites and plant hormones, transport of metabolites, signaling, and defense responses (Table S6). *LjHMGR1* and *LjSQE1* were significantly upregulated by LjbHLH32-overexpression (Table 2). Interestingly, LjbHLH32-OX hairy roots showed significantly elevated expression of genes (*AMY2*, *CYP88D4*, and *CYP88D5*) in the *AMY2* gene cluster (Tables 2 and S7; Krokida *et al*., 2013). The products of the *AMY2* gene cluster in *L. japonicus* are unknown. AMY2 produced β-amyrin and lupeol in a yeast expression system (Iturbe-Ormaetxe *et al*., 2003), and β-amyrin and dihydro-lupeol by agro-infiltration into tobacco leaves (Krokida *et al*., 2013). CYP88D4 and CYP88D5, homologs of CYP88D6 (involved in glycyrrhizin biosynthesis in *G. uralensis*), did not oxidize the tested triterpenes (Seki *et al*., 2008; Krokida *et al*., 2013). CYP71D353 showed dihydro-lupeol oxidation activity when transiently expressed in tobacco leaves (Krokida *et al*., 2013); however, the expression level of *CYP71D353* was not affected by overexpression of LjbHLH32 (Table 2).

**Table 2.** Changes in the expression of triterpene biosynthetic genes.

| Pathway | Name | Hy<br>(vs So) | SA<br>(vs SM) | LjbHLH32-OX<br>(vs GFP) | LjbHLH50-OX<br>(vs GFP) |
| --- | --- | --- | --- | --- | --- |
| TFs | *bHLH32* | **1.53** | **3.07** | **5.49** | 1.00 |
|  | *bHLH50* | **1.68** | **2.01** | **-0.76** | **3.00** |
| Precursor | *HMGR1* | **1.53** | **4.03** | **2.88** | 0.39 |
|  | *SQE1* | **1.22** | **3.02** | **1.30** | -0.11 |
| Betulinic acid | *OSC3 (LUS)* | **4.24** | **2.73** | **4.18** | **2.42** |
|  | *CYP716A51* | **3.92** | **4.02** | **3.18** | **1.41** |
| Soyasaponin<br>/Oleanolic acid | *OSC1 (bAS)* | -0.21 | **-1.17** | 0.038 | 0.38 |
| Soyasaponin | *LjCYP93E1* | -0.59 | -0.59 | -0.20 | -0.023 |
|  | *LjCYP72A61* | -0.87 | 0.56 | 0.20 | -0.11 |
|  | *CSyGT* | -0.47 | 0.28 | 0.11 | 0.27 |
|  | *DDMPT* | -0.87 | 0.79 | 0.20 | -0.21 |
| Ursolic acid | *OSC9 (aAS)* | - | -2.23 | - | - |
| Sterol | *OSC5 (CAS)* | -0.2 | -1.09 | 0.046 | 0.51 |
|  | *OSC7 (LAS)* | 0.57 | 0.62 | -0.30 | 0.43 |
| AMY2 gene cluster | *AMY2 (mixAS)* | -0.95 | -0.032 | **1.56** | **-0.82** |
|  | *CYP71D353* | 1.03 | -1.15 | 0.54 | 0.36 |
|  | *CYP88D4* | **2.14** | **4.34** | **1.43** | 0.41 |
|  | *CYP88D5* | **1.13** | **2.25** | **1.48** | 0.18 |

LjbHLH50-overexpression resulted in significant upregulation of several of the 23 TF genes selected by RNA-seq (Table 1), but those TF genes were not upregulated by LjbHLH32 overexpression (Table S6).

### LjbHLH32 shows similar trends to GubHLH3 in a transient effector-reporter analysis

Analyses of hairy roots indicated that overexpression of LjbHLH32 or LjbHLH50 upregulated BA biosynthesis directly or indirectly. To obtain insight into how they regulate BA biosynthesis, we examined the transactivation abilities of LjbHLH32 and LjbHLH50 against the promoter sequences of triterpene biosynthetic genes (Table S3) by transient effector-reporter analysis. Effector (*35S*::*bHLH*), reporter (*promoter*::*FLUC*), and reference (*35S*::*RLUC*) constructs were co-transfected into *Arabidopsis* mesophyll cell protoplasts and transactivation activities were measured as relative luciferase activities (FLUC/RLUC; Fig. 7). We used the combination of *GubHLH3* and *GuCYP93E3pro* as the positive control and confirmed a similar level of transactivation (39-fold) as a previous transient effector-reporter analysis using tobacco BY-2 cell protoplasts (Tamura *et al*., 2018). Similarly, GubHLH3 transactivated promoters of soyasaponin biosynthetic genes in *L. japonicus* except *OSC1pro* (*LjCYP93E3pro*, *LjCYP72A61pro*, *LjCSyGTpro*, and *LjDDMPTpro*) and the genes involved in precursor supply (*LjHMGR1pro* and *LjSQE1pro*). GubHLH3 weakly transactivated the promoter of *OSC5* (*cycloartenol synthase*), consistent with a previous study (Tamura *et al*., 2018). LjbHLH32 and LjbHLH50 did not transactivate *OSC3pr*o and *CYP716A51pro*. Unexpectedly, LjbHLH32 significantly transactivated the promoters of genes in the soyasaponin, precursor, and sterol pathways. Although the fold-changes induced by LjbHLH32 were generally lower than those by GubHLH3, the transactivation pattern of LjbHLH32 was highly correlated to that of GubHLH3 (PCC = 0.99). LjbHLH50 did not positively affect any promoters and the results were not correlated to those of GubHLH3 (PCC = 0.36).

**Fig. 7.**
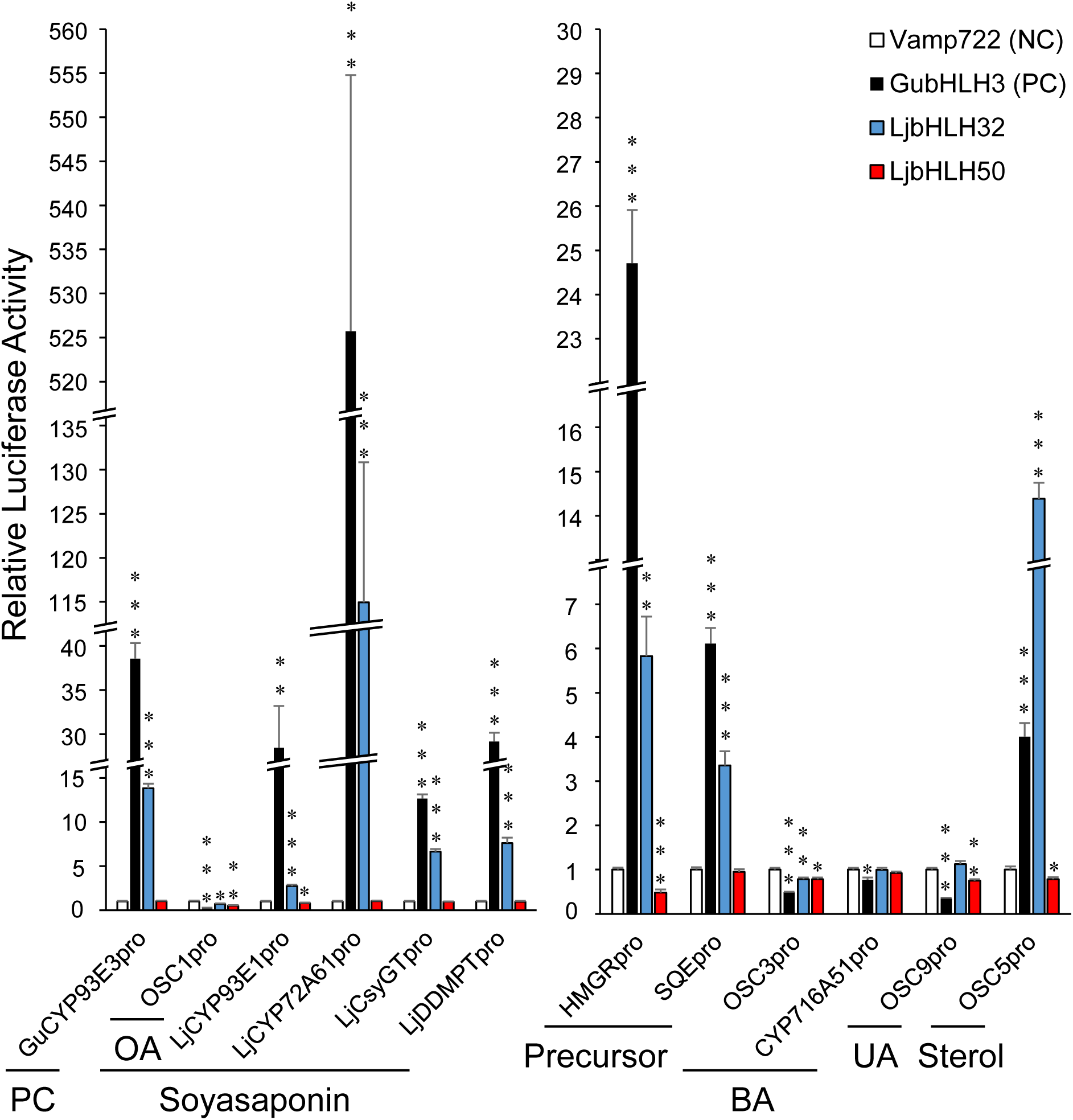
Transient effector-reporter analysis of LjbHLH32 and LjbHLH50 for the promoters of triterpene biosynthetic genes. Effector, reporter, and reference plasmids were co-transfected into *Arabidopsis* mesophyll cell protoplasts. Luciferase (LUC) activity levels were measured after overnight incubation. Error bars indicate standard errors (SE, n = 4). Asterisks indicate significant differences between the negative control (NC) and transcription factors (TFs) by Dunnett’s test (*P < 0.05, **P < 0.01, ***P < 0.001). PC, positive control; OA, oleanolic acid; BA, betulinic acid; UA, ursolic acid.

## Discussion

### SA formation and BA biosynthesis

Plant specialized metabolites are often produced and accumulated in specific cell types and organs, such as artemisinin in *Artemisia annua* (Duke & Paul, 1993; Duke *et al*., 1994), nicotine in tobacco (Katoh *et al*., 2005), glycyrrhizin in *Glycyrrhiza* spp. (Kojoma *et al*., 2010; Li *et al*., 2014), and monoterpene indole alkaloid (MIA) in *Catharanthus roseus* (Yamamoto *et al*., 2019). In this study, we identified SA as a sink tissue of BA in *L. japonicus*, which is formed on the surface of hypocotyls and roots when cultured hydroponically (Figs. 2 and S4). BA negatively regulates nodulation by controlling *ENOD40* (early nodulin gene) expression in *L. japonicus* (Delis *et al*., 2011; Suzuki *et al*., 2019). More BA accumulated in SA than in roots of soil-grown plants (Figs. 1A and 2), suggesting an unknown and SA-specific role of BA. The role of aerenchyma, including SA, is to supply air to roots under flooded conditions (Colmer & Pedersen, 2008; Shimamura *et al*., 2010; Takahashi *et al*., 2014). Our recent study revealed that *G. max* also accumulated BA in SA under flooded conditions and suggested that the accumulation of hydrophobic BA was essential to exclude water from gas spaces and promote air diffusion (Takahashi et al. 2022).

Sucrose supply from leaves is required for SA formation (Striker & Colmer, 2017; Takahashi *et al*., 2018). However, it remains elusive whether sucrose functions only as a nutrient during cell differentiation from SM to SA, or also as a signaling molecule (Tognetti *et al*., 2013). Hydroponically cultured *L. japonicus* will be useful to investigate such molecular mechanisms of SA formation.

Triterpene biosynthesis is downregulated in underground tissues of *Glycyrrhiza* spp. lacking leaves during winter (Hayashi *et al*., 1998; Ramilowski *et al*., 2013), and the total amount of triterpene decreased in aseptic liquid cultures of *G. uralensis* stolons lacking aerial parts (Kojoma *et al*., 2010; Tamura *et al*., 2017). Liquid-cultured *L. japonicus* adventitious roots also showed decreased triterpene contents compared to the roots of soil-grown plants (Fig. 1A). Photosynthesis and sucrose supply from aerial parts may affect not only SA formation but also triterpene biosynthesis in underground tissues of Fabaceae plants.

### Subclade IVa1 and IVa2 bHLHs are likely functionally diverged in Fabaceae triterpene biosynthesis regulation

Clade IVa bHLHs are conserved factors regulating jasmonate-inducible terpene metabolism in plants (Goossens *et al*., 2017; da Silva Magedans *et al*., 2020), including that of MIA in *C. roseus* (Van Moerkercke *et al*., 2015, 2016), and triterpene saponins in *Chenopodium quinoa* (Jarvis *et al*., 2017), *Panax notoginseng* (Zhang *et al*., 2017), and Fabaceae plants (Mertens *et al*., 2016; Tamura *et al*., 2018; Ribeiro *et al*., 2020). We showed that clade IVa bHLHs in Fabaceae are subdivided into three subclades; all saponin biosynthesis activators identified to date in Fabaceae fell into subclade IVa1 (Suzuki *et al*., 2021). BA biosynthesis is jasmonate-independent in Fabaceae plants (Hayashi *et al*., 2003; Broeckling *et al*., 2005; Suzuki *et al*., 2005), and LjbHLH32, a member of subclade IVa2, activated BA biosynthesis in *L. japonicus* transgenic hairy roots (Fig. 5). These results suggest that subclade IVa2 gained new target specificities during the evolution from subclade IVa1 in Fabaceae plants.

Interestingly, similar to the soyasaponin biosynthesis activator GubHLH3, LjbHLH32 transactivated the promoters of soyasaponin biosynthetic genes but not of BA biosynthetic genes in transient effector-reporter assays using *A. thaliana* mesophyll cell protoplasts (Fig. 7). The changes in *cis*-elements in promoters and DNA-binding domains in TFs are likely important to rewire gene regulatory networks (Shoji, 2019). In the case of MtTSARs, GubHLH3, and LjbHLH32, however, peptide sequences of the basic domains are well conserved (Fig. S7). Differences in protein-protein interactions among TFs may contribute to changes in target recognition by TFs. The bHLH-MYB complex is implicated in the transcriptional regulation of anthocyanin biosynthesis. In kiwifruit, *Actinidia chinensis*, the complex of AcbHLH42 and AcMYB123 regulated tissue-specific anthocyanin biosynthesis (Wang *et al*., 2019). Although single overexpression of AcbHLH42 or AcMYB123 in *Arabidopsis* was not enough to increase the anthocyanin content, their co-expression induced anthocyanin production in seedlings (Wang *et al*., 2019). LjbHLH32 may have missed its interacting partners in *Arabidopsis* mesophyll cells (Fig. 7), unlike *L. japonicus* hairy roots (Fig. 5). To prove this hypothesis, it is necessary to identify direct targets and interacting partners of LjbHLH32.

### Conserved roles of clade Ia bHLH TFs in BA biosynthesis regulation

Clade Ia bHLHs are subdivided into the SMF subclade (described in detail later) regulating stomatal development and the others subclade (Ran *et al*., 2013). BpbHLH9 in the others subclade is an OA and BA biosynthesis regulator in *B. platyphylla* (Yin *et al*., 2017). LjbHLH50 is also of the others subclade and enhanced BA biosynthesis in *Lotus* transgenic hairy roots (Fig. 5), suggesting that bHLHs in this subclade are conserved regulators of BA biosynthesis in fabids.

LjbHLH50 did not transactivate the promoters tested (Fig. 7), suggesting that it lacked its interacting partners in *Arabidopsis* mesophyll cells or indirectly regulated BA biosynthesis via downstream TF cascades. Although overexpression of BpbHLH9 enhanced expression of some triterpene biosynthesis genes and accumulation of triterpene compounds, the genes directly regulated by BpbHLH9 are unknown (Yin *et al*., 2017). Our first transcriptome analysis of whole roots from soil-grown and hydroponically cultured plants, and SA and SM cell layers isolated by LMD revealed 23 candidate TFs, including LjbHLH32 and LjbHLH50 (Fig. 4 and Table 1). We also performed transcriptome profiling of transgenic hairy root lines (Fig. 6). We found upregulation of 6 of 23 TF candidate genes by overexpression of LjbHLH50, which were annotated as LBD, ERF, NAC, and SCREAM (SCRM)-like proteins (Table S6). LjbHLH50 (and BpbHLH9) may indirectly regulate BA biosynthesis by upregulating these TF genes.

In higher plants, SPEECHLESS (SPCH), MUTE, and FAMA in the SMF subclade regulate stomatal development (Ohashi-Ito & Bergmann, 2006; MacAlister *et al*., 2007; Pillitteri *et al*., 2007). The clade IIIb bHLHs SCRM (ICE1) and SCRM2 are essential for stomatal development because they formed heterodimers with SPCH, MUTE, and FAMA (Kanaoka *et al*., 2008). AtSCRM protein (At3g26744.1; 494 aa) has bHLH (298–350 aa) and putative zipper (ZIP; 351–380 aa) domains (Chinnusamy *et al*., 2003). Alignments between AtSCRM and AtSCRM-like (At2g40435.1) proteins showed that SCRM-like proteins lacked a bHLH domain but possessed a partial ZIP domain (Fig. S8). An analysis of the interaction between LjbHLH50 (BpbHLH9) and SCRM-like proteins is interesting.

### Sleeping BA biosynthesis ability in *M. truncatula*

The *M. truncatula* genome has putative lupeol synthase genes but lupeol derivatives have not been detected in *Medicago* spp. (Naoumkina *et al*., 2010). We found that whole roots from hydroponically cultured plants and liquid-cultured adventitious roots accumulated trace amounts of BA (Fig. 1C). Because homologs of LjbHLH32 and LjbHLH50 were present in *M. truncatula* (Fig. 4D), expression of these TFs might limit BA biosynthesis in this plant.

Silencing of *OSC3* expression and reduced accumulation of lupeol derivatives results in upregulated *ENOD40* expression and enhanced nodulation in *L. japonicus* (Delis *et al*., 2011). Similarly, in *M. truncatula* roots, overexpression of chimeric MtbHLH1 fused to the EAR repression domain (transgenic plants should show dominant loss-of-function phenotypes; Hiratsu *et al*., 2003) resulted in abnormal nodule vascular bundle development and slightly increased numbers of nodules per plant although the growth of untransformed aerial parts was significantly reduced in the composite plants (Godiard *et al*., 2011). Although it is unknown whether MtbHLH1 positively regulates BA biosynthesis similar to its ortholog LjbHLH50 and whether *M. truncatula* nodules accumulate lupeol derivatives, Delis *et al*. (2011) and Godiard *et al*. (2011) likely reported similar phenomena. Re-evaluation of these mutants will provide rigid understanding for the role of lupeol derivatives and clade Ia bHLHs in nodule formation.

Godiard *et al*. (2011) suggested that an LBD-type TF gene (Medtr7g007010) is a potential target of MtbHLH1 by transcriptome analysis of transgenic roots overexpressing MtbHLH1-EAR. An LBD gene was co-upregulated in hydroponic roots, SA, and LjbHLH50-OX hairy roots (Lj5g3v2099760/LotjaGi5g1v0323400; Table S6) and closely related to Medtr07g007010 in the phylogenetic tree (Fig. S9), suggesting the involvement of LBD proteins in this yet-to-be-defined TF cascade.

### Concluding remarks

We identified two bHLH-type TFs enhancing BA biosynthesis by transcriptome analysis of hydroponically cultured *L. japonicus* and a functional analysis using transgenic hairy roots. However, neither LjbHLH32 nor LjbHLH50 transactivated the promoters of BA biosynthetic genes in transient effector-reporter assays using a heterologous host (*Arabidopsis*), suggesting that these bHLHs interact with unknown TFs or indirectly regulate BA biosynthesis via downstream TFs. The 23 putative TFs found in the first screening are promising targets for further analysis (Table 1). This study is a first step toward understanding the complicated transcriptional regulatory mechanisms of BA biosynthesis in *L. japonicus*.

## Supplementary data

**Fig. S1.** Transfer DNA regions of gateway destination vectors for *L. japonicus* transformation.

**Fig. S2.** Triterpene biosynthetic pathways in *L. japonicus*.

**Fig. S3.** Mass spectra of peaks 6 and 10 in the chromatograms in Fig. 1.

**Fig. S4.** Time-course-dependent increase in BA in hydroponic roots of *L. japonicus*.

**Fig. S5.** Magnified image of secondary aerenchyma from a hydroponic *L. japonicus* plant.

**Fig. S6.** MA plots for differentially expressed gene (DEG) detection.

**Fig. S7.** Protein alignments of the basic domains of clade IVa bHLHs.

**Fig. S8.** Alignments of AtSCRM and AtSCRM-like proteins.

**Fig. S9.** Phylogenetic tree of LBD (LOB) proteins in *M. truncatula* and *L. japonicus*.

**Table S1.** List of primers.

**Table S2.** List of FastQ files obtained.

**Table S3.** Promoter sequences isolated in this study.

**Table S4.** List of differentially expressed genes (DEGs) identified in this study.

**Table S5.** Annotation of 256 differentially expressed genes (DEGs) commonly upregulated in Hy and SA samples.

**Table S6.** Annotation of differentially expressed genes (DEGs) in Fig. 6C.

**Table S7.** Detailed results related to triterpene biosynthesis genes obtained by TCC-GUI analysis.

## Acknowledgements

We thank Miyazaki University and National BioResource Project (NBRP) for providing pUB-GWS-GFP plasmid and *L. japonicus* Gifu B-129 seeds, RIKEN BRC for providing Lj suspension cell line (rpc00032) through NBRP, the Institut National de la Recherche Agronomique for providing *M. truncatula R108* and Jemalong A17 seeds, Prof. Shuhei Yasumoto (Osaka University) for providing genomic DNA of *A. thaliana* Col-0, and Dr. Atsushi Toyoda (National Institute of Genetics) for construction and sequencing of RNA-seq library of soil-grown and hydroponically-cultured whole roots. We also thank Dr. Nobutaka Mitsuda, Dr. Shingo Sakamoto and Ms. Tomoko Niki (AIST) for technical support on transient effector-reporter analysis.

## Author Contribution

EOF, TM and HSe conceived and supervised the study; All authors designed the experiments; HSu and HT performed the experiments; HSu wrote the manuscript; HT, EOF, MN, TM and HSe made manuscript revisions. All authors read and approved the final manuscript.

## Data Availability

The nucleotide sequence reported in this paper has been submitted to the DNA Data Bank of Japan (DDBJ) under the accession numbers of LC609881 (*LjbHLH32*) and LC609882 (*LjbHLH50*). The raw RNA-seq reads obtained were deposited in the DDBJ Sequence Read Archive (DRA) under the accession number DRA011635.

The English in this document has been checked by at least two professional editors, both native speakers of English. For a certificate, please see: http://www.textcheck.com/certificate/3bPF0g

