## Supplementary material for "Identification of basic helix-loop-helix transcription factors that activate betulinic acid biosynthesis by RNA-sequencing of hydroponically cultured *Lotus japonicus*": Fig. S

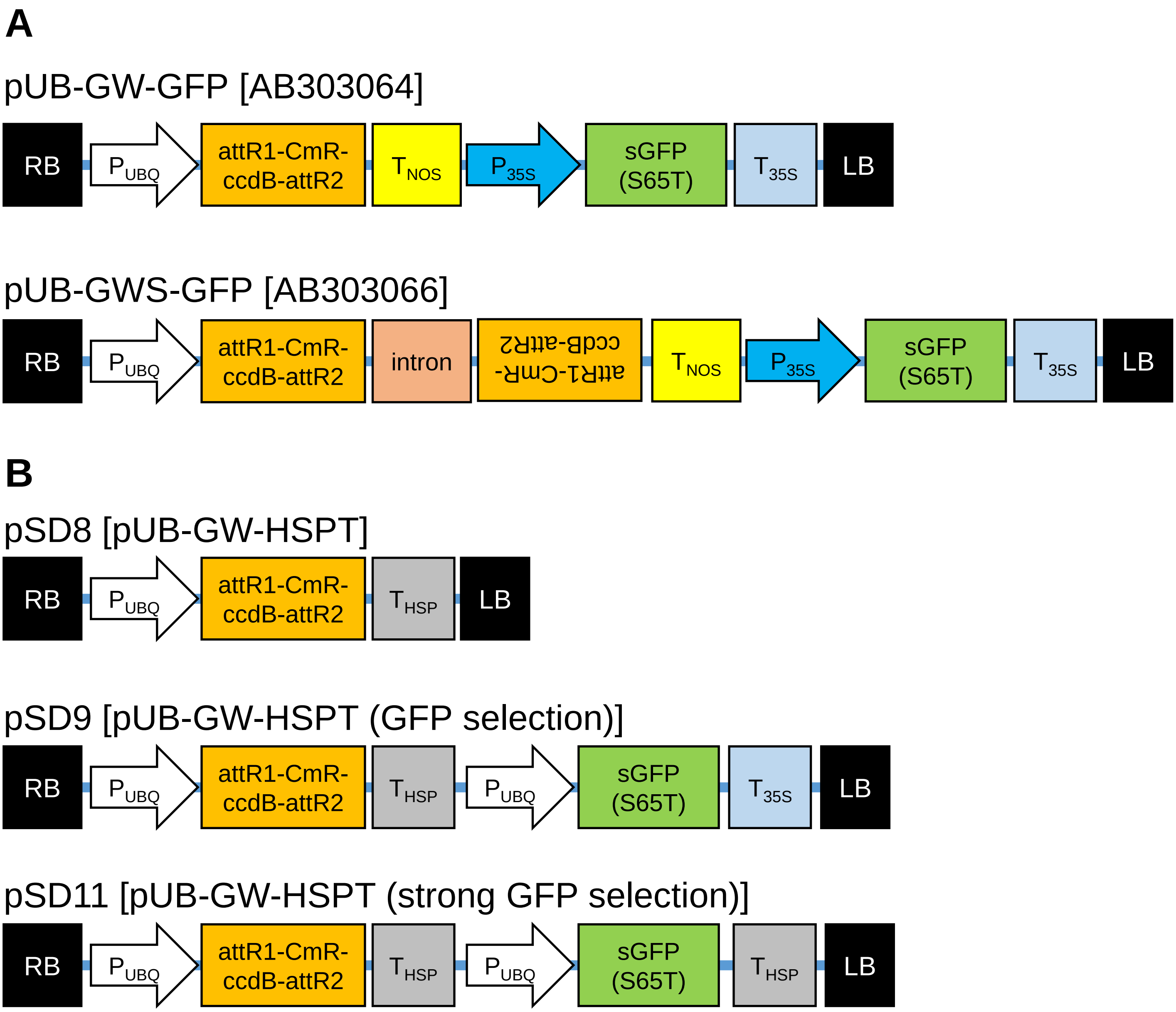


**Fig. S1. Transfer DNA regions of gateway destination vectors for *L. japonicus* transformation.** (A) Vectors provided by the National Bioresource Project. (B) New vectors created from the pUB-GWS-GFP vector. pSD11 was used for hairy root transformation to overexpress TF genes. $P_{UBQ}$, *L. japonicus* polyubiquitin (*LjUBQ1*) promoter; $P_{35S}$, cauliflower mosaic virus 35S promoter; $T_{35S}$, 35S terminator; $T_{NOS}$, nopaline synthase terminator; $T_{HSP}$, terminator of *Arabidopsis thaliana* heat shock protein 18.2 (HSP18.2) gene; intron, intron1 of *AtWRKY33*; CmR, chloramphenicol resistance gene; ccdB, counter-selectable marker gene; RB, right border; LB, left border.

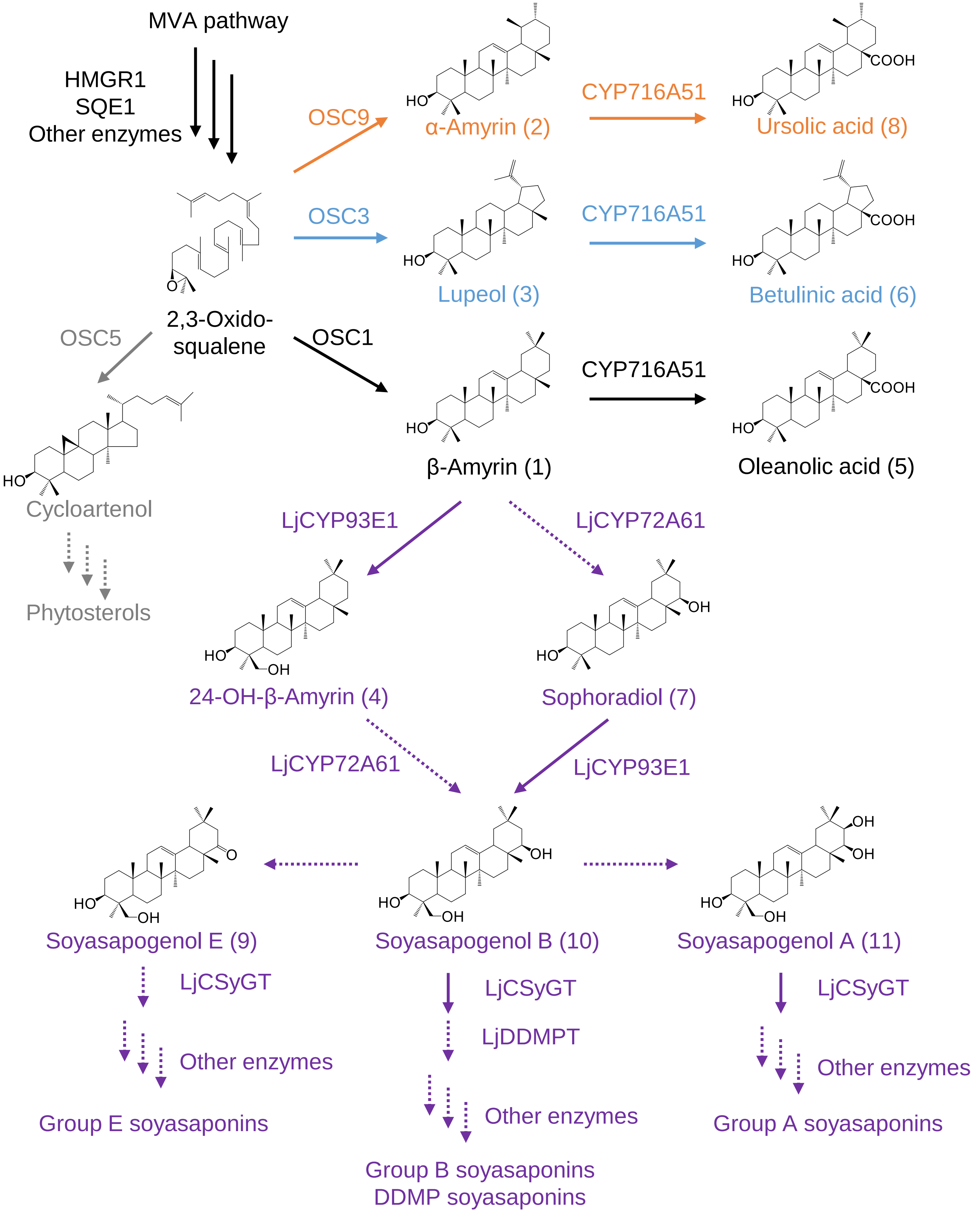


**Fig. S2. Triterpene biosynthetic pathways in *L. japonicus*.** Dashed arrows indicate the reactions putatively catalyzed by the enzymes. MVA, mevalonate; HMGR, 3-hydroxy-3-methylglutaryl-CoA reductase; SQE, squalene epoxidase; OSC, oxidosqualene cyclases; CYP, cytochrome P450 monooxygenases; CSyGT, cellulose-synthase-derived glycosyltransferase; DDMPT, 2,3-dihydro-3,5-dihydroxy-6-methyl-4H-pyran-4-one transferase.

Peak 6: Betulinic acid (BA)

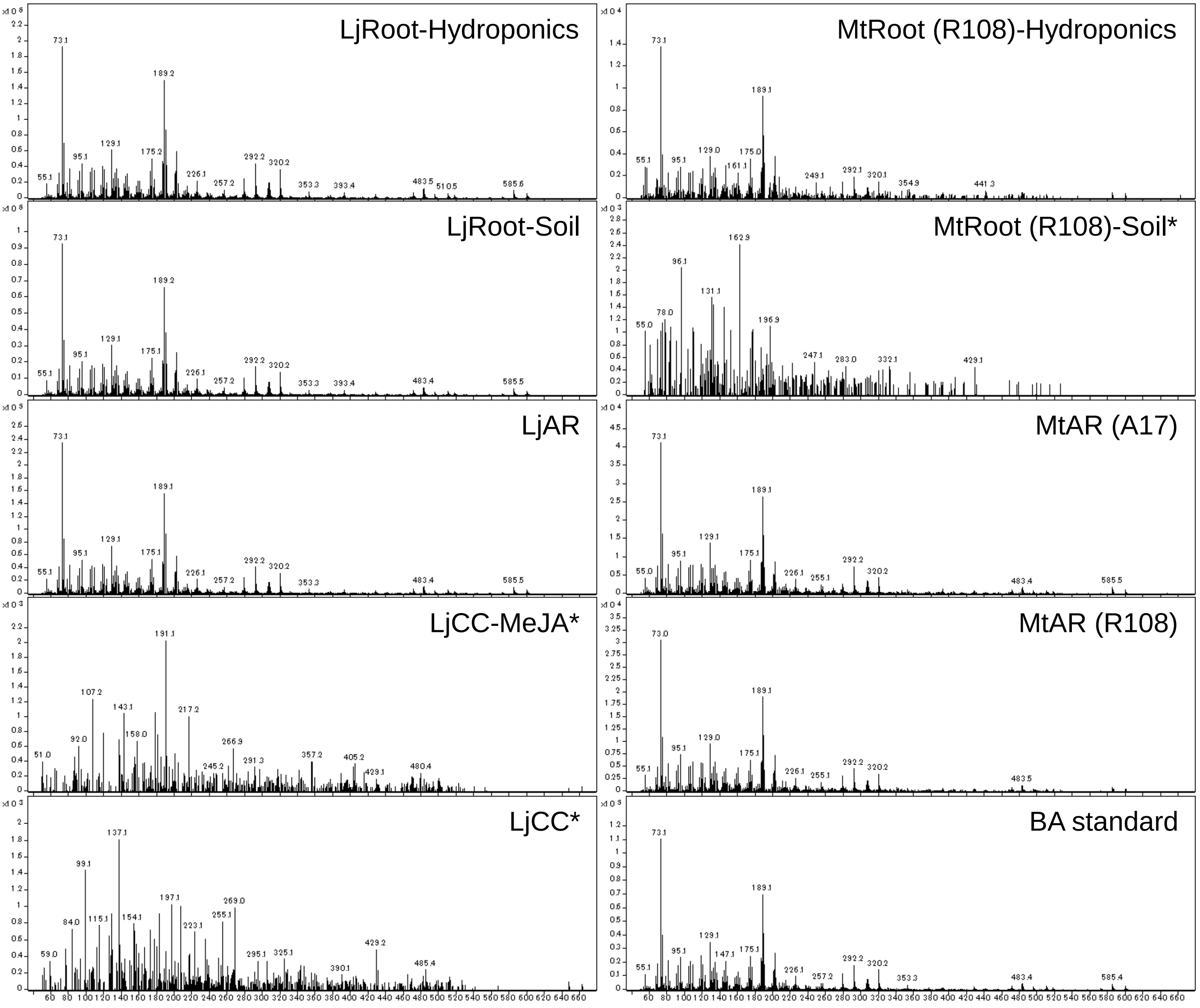


Peak 10: Soyasapogenol B (SB)

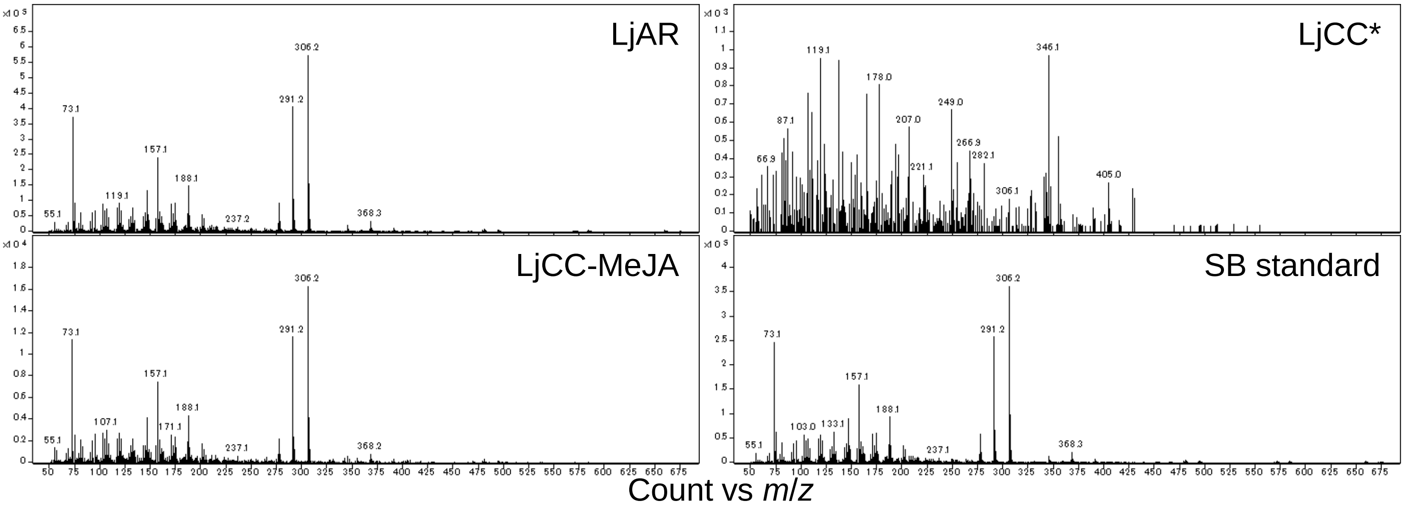


**Fig. S3. Mass spectra of peaks 6 and 10 in the chromatograms in Fig. 1.** *The mass fragmentation patterns were not consistent with the authentic standard compounds. Lj, *Lotus japonicus*; Mt, *Medicago truncatula*; AR, adventitious roots; CC, cultured cells; CC-MeJA, cultured cells treated with methyl jasmonate; A17, cultivar Jemalong A17.

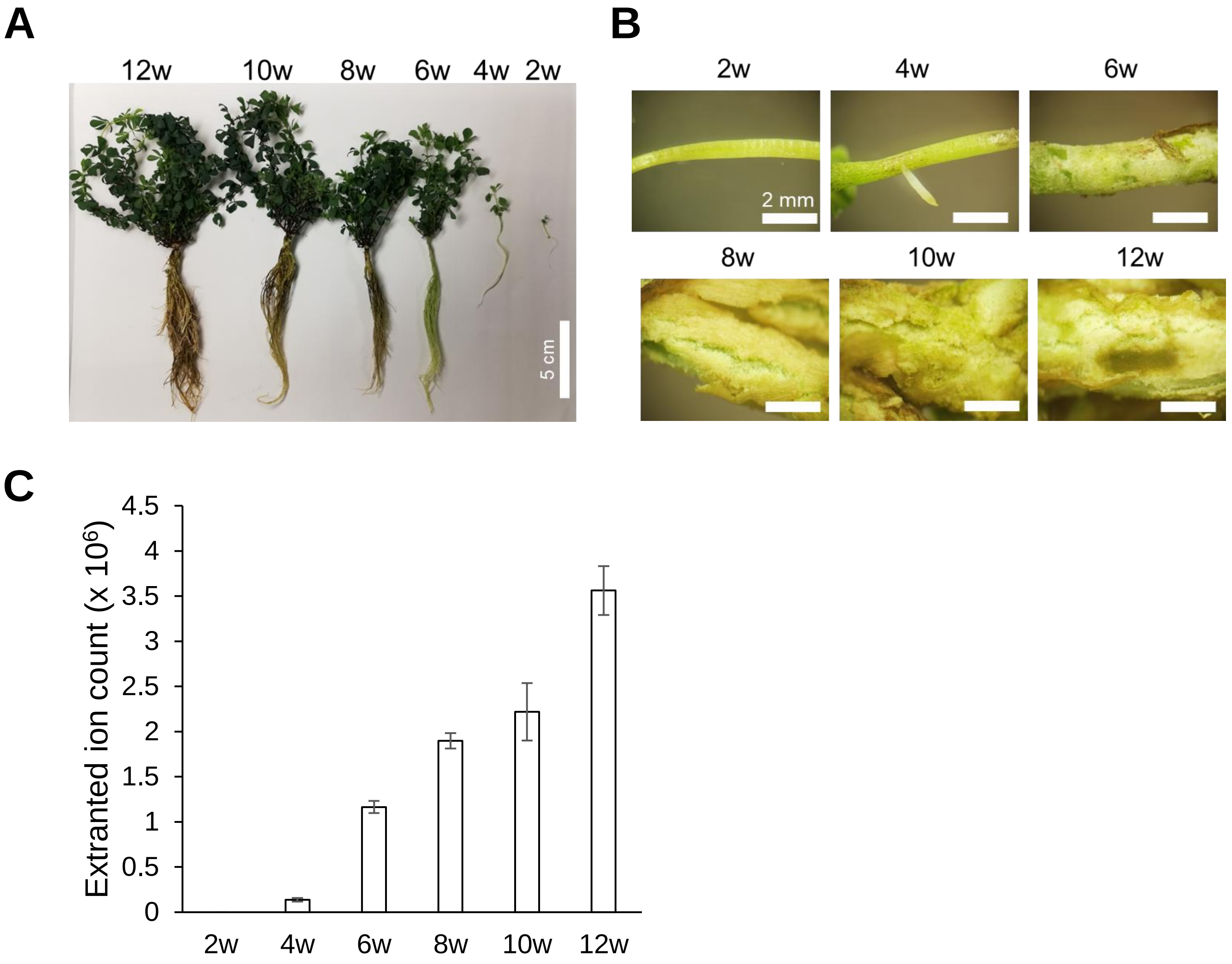


**Fig. S4. Time-course-dependent increase in BA in hydroponic roots of *L. japonicus*.** Whole plants (A) and hypocotyls (B) of 2–12-week-old seedlings cultured hydroponically. (C) The peak area of BA was calculated from the extracted ion counts (*m/z* = 189).

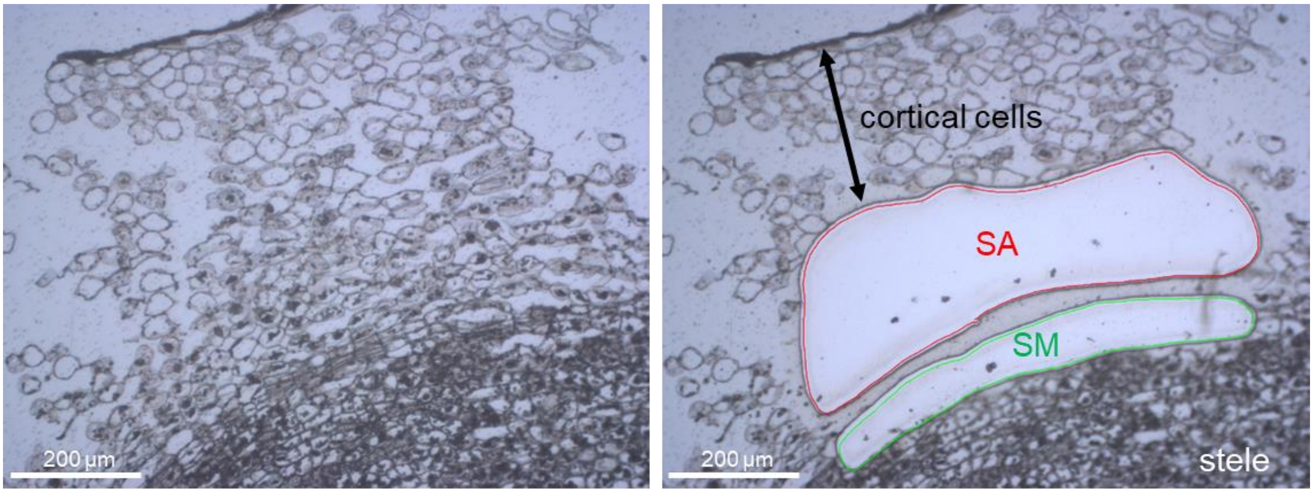


**Fig. S5. Magnified image of secondary aerenchyma from a hydroponic *L. japonicus* plant.** Secondary aerenchyma (SA) and secondary meristem (SM) were isolated by laser microdissection (LMD).

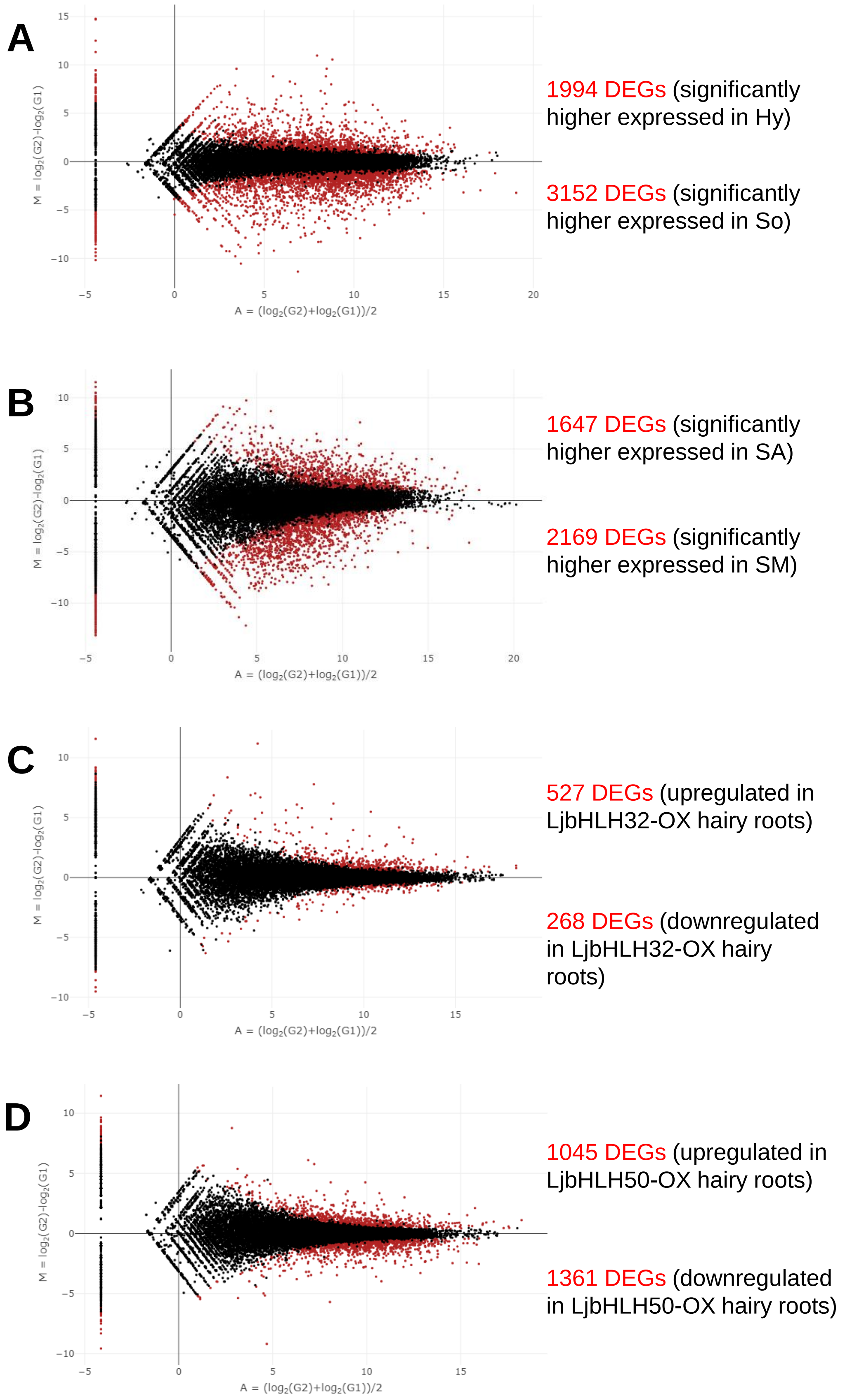


**Fig. S6. MA plots for differentially expressed gene (DEG) detection.** Differential expression analysis was performed between two groups using the graphical user interface for TCC (TCC-GUI). Group 1 (G1) and G2 are soil-cultured whole roots (So) and hydroponic whole roots (Hy) **(A)**, secondary meristem (SM) and secondary aerenchyma (SA) **(B)**, GFP-expressing and LjbHLH32-OX hairy roots **(C)**, and GFP-expressing and LjbHLH50-OX hairy roots **(D)**, respectively. M value, $\log_2$ (fold change) indicates the difference of expression level between groups. a.value, $\log_2$ (mean expression level). DEGs are indicated in red. The threshold of the false discovery rate (FDR) is $\leq 0.10$.

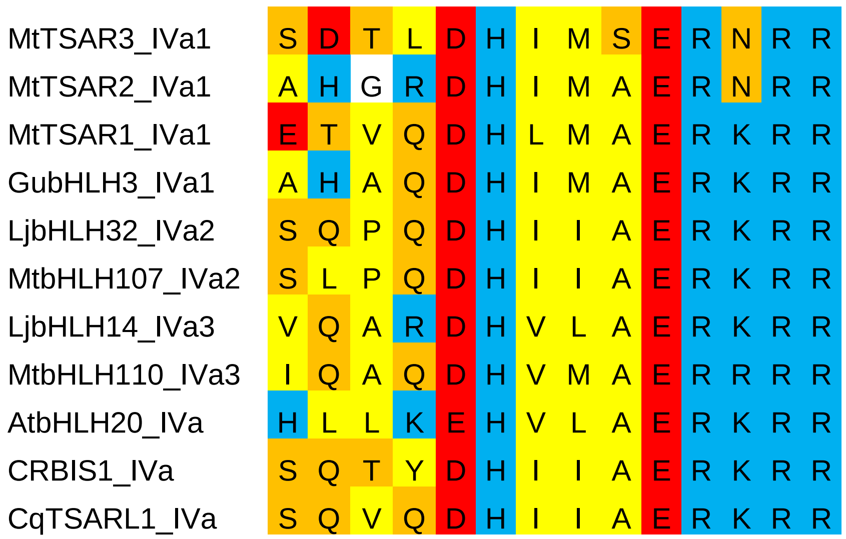


**Fig. S7. Protein alignments of the basic domains of clade IVa bHLHs.** Basic, acidic, hydrophobic, and hydrophilic amino acids are shown as a blue, red, yellow, and orange background, respectively.

```
Query: AtSCRM1_At3g26744.1
Sbjct: AtSCRM-like_At2g40435.1
Identities:44/187(24%), Positives:88/187(47%), Gaps:35/187(18%)

Query  309  LMAERRRRKKLNDRLYMLRSVVPKISKMDRASILGDAIDYLKELLQRINDLHNELESTPP  368
            +++  ++R  L ++  +LRS+     ++D  SI+ DA  Y+++L Q++   + +
Sbjct  1    MVSREQKRGSLQEKFQLLRSITNSHAEND-TSIIMDASKYIQKLKQKVERFNQD------  53

Query  369  GSLPPTSSSFHPLTPTPQTLSCRVKEELCPSSLPSPKGQQARVEVRLREGRAVNIHMFCG  428
                PT+                   E   S    PK     VE  L +G  +N+  F G
Sbjct  54   ----PTA-------------------EQSSSEPTDPKTPMVTVET-LDKGFMINV--FSG  87

Query  429  R-RPGLLLATMKALDNLGLDVQQAVISCFNGFALDVFRAEQCQEGQEILPDQIKAVLFDT  487
            + +PG+L++ ++A +++GL+V +A  SC + F+L     E  ++G+ +  + +K  + D
Sbjct  88   KNQPGMLVSVLEAFEDIGLNVLEARASCTDSFSLHAMGLEN-EDGENMDAEAVKQAVTDA  146

Query  488  AGYAGMI  494
                G I
Sbjct  147  IRSWGEI  153
```

**Fig. S8. Alignments of AtSCRM and AtSCRM-like proteins.**

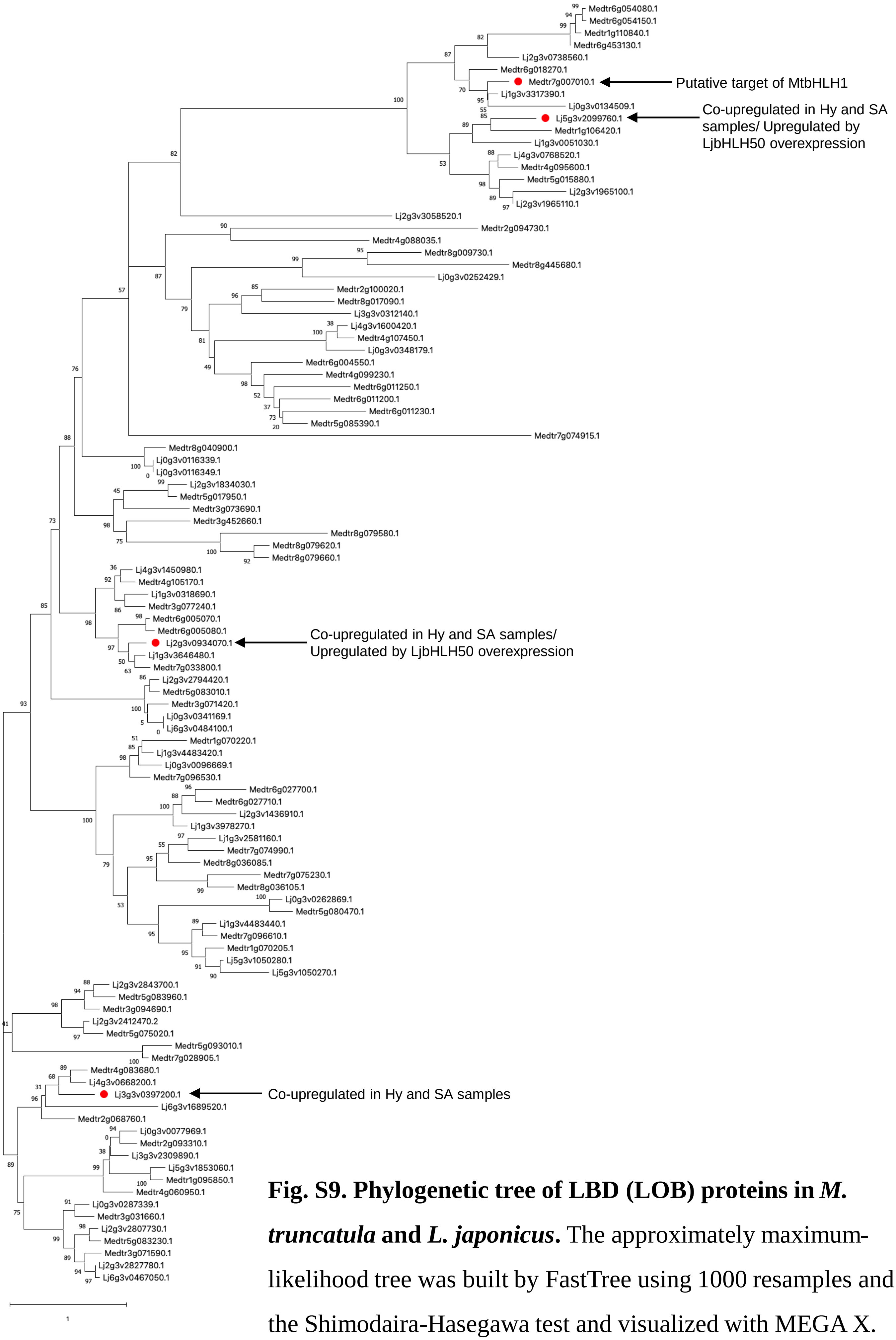


**Fig. S9. Phylogenetic tree of LBD (LOB) proteins in *M. truncatula* and *L. japonicus*.** The approximately maximum-likelihood tree was built by FastTree using 1000 resamples and the Shimodaira-Hasegawa test and visualized with MEGA X.
